# Quantifying the impact of an inference model in Bayesian phylogenetics

**DOI:** 10.1101/2019.12.17.879098

**Authors:** Richèl J.C. Bilderbeek, Giovanni Laudanno, Rampal S. Etienne

**Affiliations:** Groningen Institute for Evolutionary Life Sciences, University of Groningen, Groningen, The Netherlands

**Keywords:** Bayesian model selection, BEAST2, computational biology, evolution, phylogenetics, R, tree prior, babette

## Abstract

1. Phylogenetic trees are currently routinely reconstructed from an alignment of character sequences (usually nucleotide sequences). Bayesian tools, such as MrBayes, RevBayes and BEAST2, have gained much popularity over the last decade, as they allow joint estimation of the posterior distribution of the phylogenetic trees and the parameters of the underlying inference model. An important ingredient of these Bayesian approaches is the species tree prior. In principle, the Bayesian framework allows for comparing different tree priors, which may elucidate the macroevolutionary processes underlying the species tree. In practice, however, only macroevolutionary models that allow for fast computation of the prior probability are used. The question is how accurate the tree estimation is when the real macroevolutionary processes are substantially different from those assumed in the tree prior.
2. Here we present `pirouette`, a free and open-source R package that assesses the inference error made by Bayesian phylogenetics for a given macroevolutionary diversification model. `pirouette` makes use of BEAST2, but its philosophy applies to any Bayesian phylogenetic inference tool.
3. We describe `pirouette`’s usage providing full examples in which we interrogate a model for its power to describe another.
4. Last, we discuss the results obtained by the examples and their interpretation.

## 1 Introduction

The development of new powerful Bayesian phylogenetic inference tools, such as BEAST [Drummond & Rambaut 2007], MrBayes [Huelsenbeck & Ronquist 2001] or RevBayes [Höhna *et al*. 2016a] has been a major advance in constructing phylogenetic trees from character data (usually nucleotide sequences) extracted from organisms (usually extant, but extinction events and/or time-stamped data can also be added), and hence in our understanding of the main drivers and modes of diversification.

BEAST [Drummond & Rambaut 2007] is a typical Bayesian phylogenetics tool, that needs both character data and priors to infer a posterior distribution of phylogenies. Specifically, for the species tree prior - which describes the process of diversification - BEAST has built-in priors such as the Yule [Yule 1925] and (constant-rate) birth-death (BD) [Nee *et al*. 1994] models as well as coalescent priors. These simple tree priors are among the most commonly used, as they represent some biologically realistic processes (e.g. viewing diversification as a branching process), while being computationally fast.

To allow users to extend the functionalities of BEAST using plug-ins, BEAST2 was written [Bouckaert *et al*. 2019] (with BEAST and BEAST2 still independently being developed further). For example, one can add novel diversification models by writing a BEAST2 plugin that contains the likelihood formula of a phylogeny under the novel diversification model, i.e. the prior probability of a species tree. Plugins have been provided, for instance, for the calibrated Yule model [Heled & Drummond 2015], the BD model with incomplete sampling [Stadler 2009], the BD model with serial sampling [Stadler *et al*. 2012], the BD serial skyline model [Stadler *et al*. 2013], the fossilized BD process [Gavryushkina *et al*. 2014], and the BD SIR model [Kühnert *et al*. 2014].

Many other diversification models (and their associated likelihood algorithms) have been developed, e.g., models in which diversification is time-dependent [Nee *et al*. 1994; Rabosky & Lovette 2008], or diversity-dependent [Etienne *et al*. 2012], or where diversification rates change for specific lineages and their descendants [Etienne & Haegeman 2012; Rabosky 2014; Alfaro *et al*. 2009; Laudanno *et al*. 2020]. Other models treat speciation as a process that takes time [Rosindell *et al*. 2010; Etienne & Rosindell 2012; Lambert *et al*. 2015], or where diversification rates depends on one or more traits [Maddison *et al*. 2007; FitzJohn 2012; Herrera-Alsina *et al*. 2019].

These are, however, not yet available as tree priors in BEAST2, for reasons explained below. In this paper, we present methodology to determine whether such new plug-ins are needed, or whether currently available plug-ins are sufficient. We show this using the Yule and BD species tree priors, but our methods can be used with other built-in tree priors as well.

The rationale of our paper is as follows. When a novel diversification model is introduced, its performance in inference should be tested. Part of a model’s performance is its ability to recover parameters from simulated data with known parameters (e.g. [Etienne *et al*. 2014]), where ideally the estimated parameter values closely match the known/true values. Even when a diversification model passes this test, it is not necessarily used as tree prior in Bayesian inference. Bayesian phylogenetic inference often requires that the prior probability of the phylogeny according to the diversification model has to be computed millions of times. Therefore, biologically interesting but computationally expensive tree priors are often not implemented, and simpler priors are used instead. This is not necessarily problematic, when the data are very informative or when the prior is truly uninformative, as this will reduce the influence of the tree prior. However, the assumption that tree prior choice is of low impact must first be verified.

There have been multiple attempts to investigate the impact of tree prior choice. For example, Sarver and colleagues, [Sarver *et al*. 2019] showed that the choice of tree prior does not substantially affect phylogenetic inferences of diversification rates. However, they only compared current diversification models to one another, and thus this does not inform us on the impact of a new tree prior.

Similarly, Ritchie and colleagues [Ritchie *et al*. 2016] showed that inference was accurate when birth-death or skyline coalescent priors were used, but they simulated their trees with a Yule process only, as their focus was not so much on the diversification process but on the influence of inter- and intraspecific sampling.

Another way to benchmark a diversification model, is by doing a model comparison, in which the best model is determined from a set of models. A good early example is Goldman 1993 in which Goldman compared DNA substitution models. A recent approach to test the impact of tree prior choice, proposed by Duchene *et al*. 2018, allows to measure model adequacy for phylodynamic models that are mathematically described (i.e. have a known likelihood equation).

Here we introduce a method to quantify the impact of a novel tree prior, i.e., a tree model, for which we can simulate phylogenies, but not yet calculate their likelihoods. This new method simultaneously assesses the substitution, clock and tree models [Duchêne *et al*. 2015]. The method starts with a phylogeny generated by the new model. Next, nucleotide sequences are simulated that follow the evolutionary history of the given phylogeny. Then, using BEAST2’s built-in tree priors, a Bayesian posterior distribution of phylogenies is inferred. We then compare the inferred with the original phylogenies. How to properly perform this comparison forms the heart of our method. Only new diversification models that result in a large discrepancy between inferred and simulated phylogenies will be worth the effort and computational burden to implement a species tree prior for in a Bayesian framework.

Our method is programmed as an R package [R Core Team 2013] called `pirouette`. `pirouette` is built on `babette` [Bilderbeek & Etienne 2018], which calls BEAST2 [Bouckaert *et al*. 2019].

## 2 Description

The goal of `pirouette` is to quantify the impact of a new tree prior. It does so by measuring the inference error made for a given reconstructed phylogeny, simulated under a (usually novel) diversification model. We refer to the model that has generated the given tree as the ‘generative tree model’ *p*_*G*_. A ‘generative tree model’, in this paper, can be either the novel diversification model for which we are testing the impact of choosing standard tree priors for, or it is the model with which we generate the twin tree that is needed for comparison (see below). In the latter case, we also refer to it as the actual generative tree model, and it thus serves as a baseline model. This is is done in the example, where the Yule model is the generative model.

The inference error we aim to quantify is not of stochastic nature. Stochastic errors are usually non-directional. We, instead, aim to expose the bias due to the mismatch between a generative model (that has generated the phylogeny) and the model(s) used in the actual inference. We define the birth-death (BD) model [Nee *et al*. 1994] as the standard tree model, as many (non-standard) tree models have a parameter setting such that it reduces to this model. One such example is the diversity-dependent (DD) diversification model [Etienne & Haegeman 2020; Etienne *et al*. 2012] in which speciation or extinction rate depends on the number of species and a clade-level carrying capacity. The BD model can be seen as a special case of the DD model, because for an infinite carrying capacity, the DD model reduces to the BD model. When benchmarking a novel tree model, one will typically construct phylogenies for different combinations of the diversification model’s parameters, to assess under which scenarios the inference error cannot be neglected. While we recommend many replicate simulations when assessing a novel tree prior, our example contains only one replicate, as the goal is to show the workings of `pirouette`, instead of doing an extensive analysis. The supplementary material includes results of replicated runs under multiple settings.

`pirouette` allows the user to specify a wide variety of custom settings. These settings can be grouped in macro-sections, according to how they operate in the pipeline. We summarize them in Table 1 and Table 2.

**Table 1:**
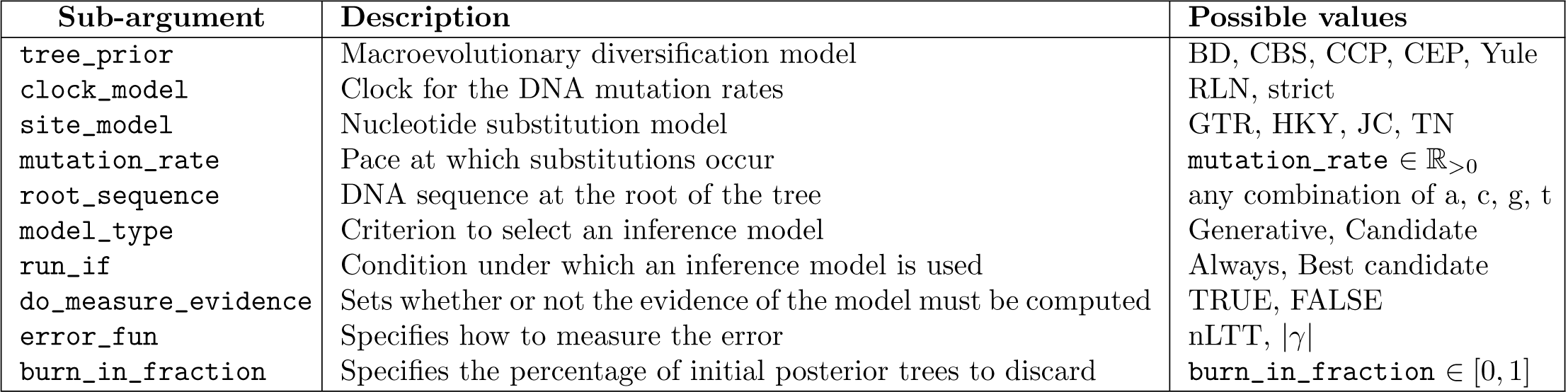
Most important parameter options. BD = birth death [Nee *et al*. 1994], CBS = coalescent Bayesian skyline [Drummond *et al*. 2005], CCP = coalescent constant-population, CEP = coalescent exponential-population, Yule = pure birth model [Yule 1925], RLN = relaxed log-normal clock model [Drummond *et al*. 2006], strict = strict clock model [Zuckerkandl & Pauling 1965], GTR = Generalized time-reversible model [Tavaré 1986], HKY = Hasegawa, Kishino and Yano [Hasegawa *et al*. 1985], JC = Jukes and Cantor [Jukes *et al*. 1969], TN = Tamura and Nei [Tamura & Nei 1993], nLTT = normalized lineages-through-time [Janzen *et al*. 2015], |*γ*| = absolute value of the gamma statistic [Pybus & Harvey 2000].

**Table 2:** Definitions of terms and relative symbols used in the main text and in Fig 1. To run the pipeline *A, X* and *E* must be specified.

| Symbol | Macro-argument | Description |
| --- | --- | --- |
| $G$ | Generative model | The full setting to produce BEAST2 input data. Its core features are the tree prior $p_G$, the clock model $c_G$ and the site model $s_G$. |
| $s_G$ | Site model | Both the substitution model and rate variation across sites. |
| $A$ | Alignment model | Specifies the alignment generation, such as the clock model $c_G$, site model $s_G$ and root sequence. |
| $X_i$ | $i$-th candidate experiment | Full setting for a Bayesian inference. It is made by a candidate inference model $I_i$ and its inference conditions $C_i$. |
| $I$ | Inference model | The assumed phylogenetic inference model, of which the main components are the tree prior $p_I$, assumed clock model $c_I$ and assumed site model $s_I$. |
| $C$ | Inference conditions | Conditions under which $I$ is used in the inference. They are composed of the model type, run condition and whether to measure the evidence. |
| $E$ | Error measure parameters | Errors measurement setup that can be specified providing an error function to measure the difference between the original phylogeny and the inferred posterior. The first iterations of the MCMC chain of the posterior may not be representative and can be discarded using a burn-in fraction. |

### 2.1 pirouette’s pipeline

The pipeline to assess the error BEAST2 makes in inferring this phylogeny contains the following steps:

1. The user supplies one or (ideally) more phylogenies from a new diversification model.
2. From the given phylogeny an alignment is simulated under a known alignment model *A*.
3. From this alignment, according to the specified inference conditions *C*, an inference model *I* is chosen (which may or may not differ from the model that generated the tree).
4. The inference model and the alignment are used to infer a posterior distribution of phylogenies.
5. The phylogenies in the posterior are compared with the given phylogeny to estimate the error made, according to the error measure *E* specified by the user.

The pipeline is visualized in Fig. 1. There is also the option to generate a ‘twin tree’, that goes through the same pipeline (see supplementary subsection 5.5).

**Figure 1:**
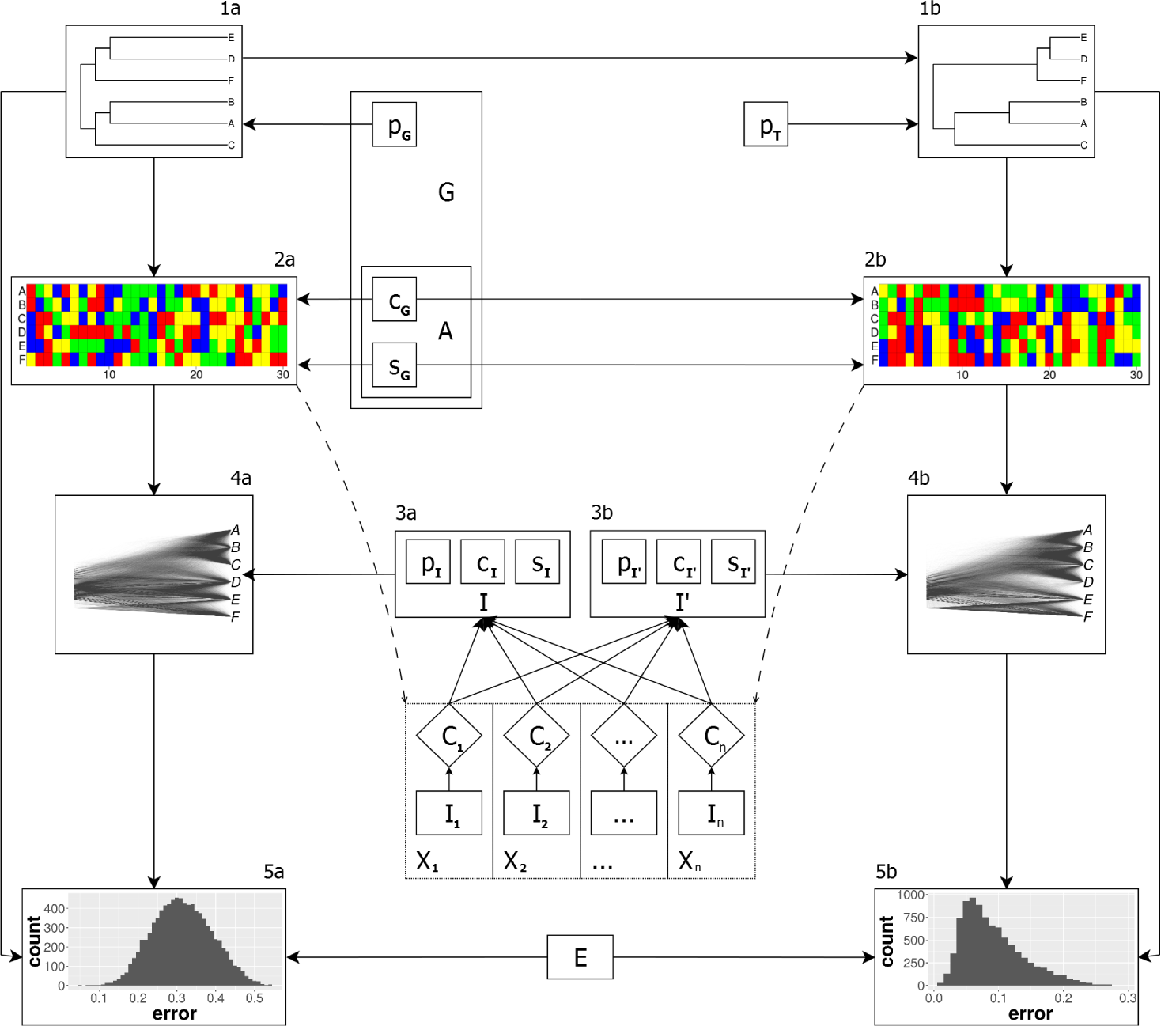
`pirouette` pipeline. The pipeline starts from a phylogeny (1a) simulated by the generative tree model *p*_*G*_. The phylogeny is converted to an alignment (2a) using the generative alignment model *A* = (*c*_*G*_, *s*_*G*_), composed of a clock model and a site model. The user defines one or more experiments. For each candidate experiment *X*_*i*_ (a combination of inference model *I*_*i*_ and condition *C*_*i*_), if its condition *C*_*i*_ is satisfied (which can depend on the alignment), the corresponding inference model *I* = *I*_*i*_ is selected to be used in the next step. The inference models (3a) of the selected experiments use the alignment (2a) to each create a Bayesian posterior of (parameter estimates and) phylogenies (4a). Each of the posterior trees is compared to the true phylogeny (1a) using the error measure *E*, resulting in an error distribution (5a). Optionally, for each selected inference model a twin pipeline can be run. A twin phylogeny (1b) can be generated from the original phylogeny (1a) using the twin tree model *p*_*t*_, selected among standard diversification models; the default option is the standard BD model, with parameters estimated from the original phylogeny. A twin alignment (2b) is then simulated from the twin phylogeny using clock model *c*_*G*_ and site model *s*_*G*_ used with the generative tree model (the novel tree model). The twin pipeline follows the procedure of the main pipeline, resulting in a twin error distribution (5b).

The first step simulates an alignment from the given phylogeny (Fig. 1, 1a → 2a). For the sake of clarity, here we will assume the alignment consists of DNA sequences, but one can also use other heritable materials such as amino acids. The user must specify a root sequence (i.e. the DNA sequence of the shared common ancestor of all species), a mutation rate and a site model.

The second step (Fig. 1, 3a) selects one or more inference model(s) *I* from a set of standard inference models *I*_1_, …, *I*_*n*_. For example, if the generative model is known and standard (which it is for the twin tree, see below), one can specify the inference model to be the same as the generative model. If the tree model is unknown or non-standard - which is the primary motivation for this paper -, one can pick a standard inference model which is considered to be closest to the true tree model. Alternatively, if we want to run only the inference model that fits best to an alignment from a set of candidates (regardless of whether these generated the alignments), one can specify these inference models (see section 5.6).

**Figure 2:**
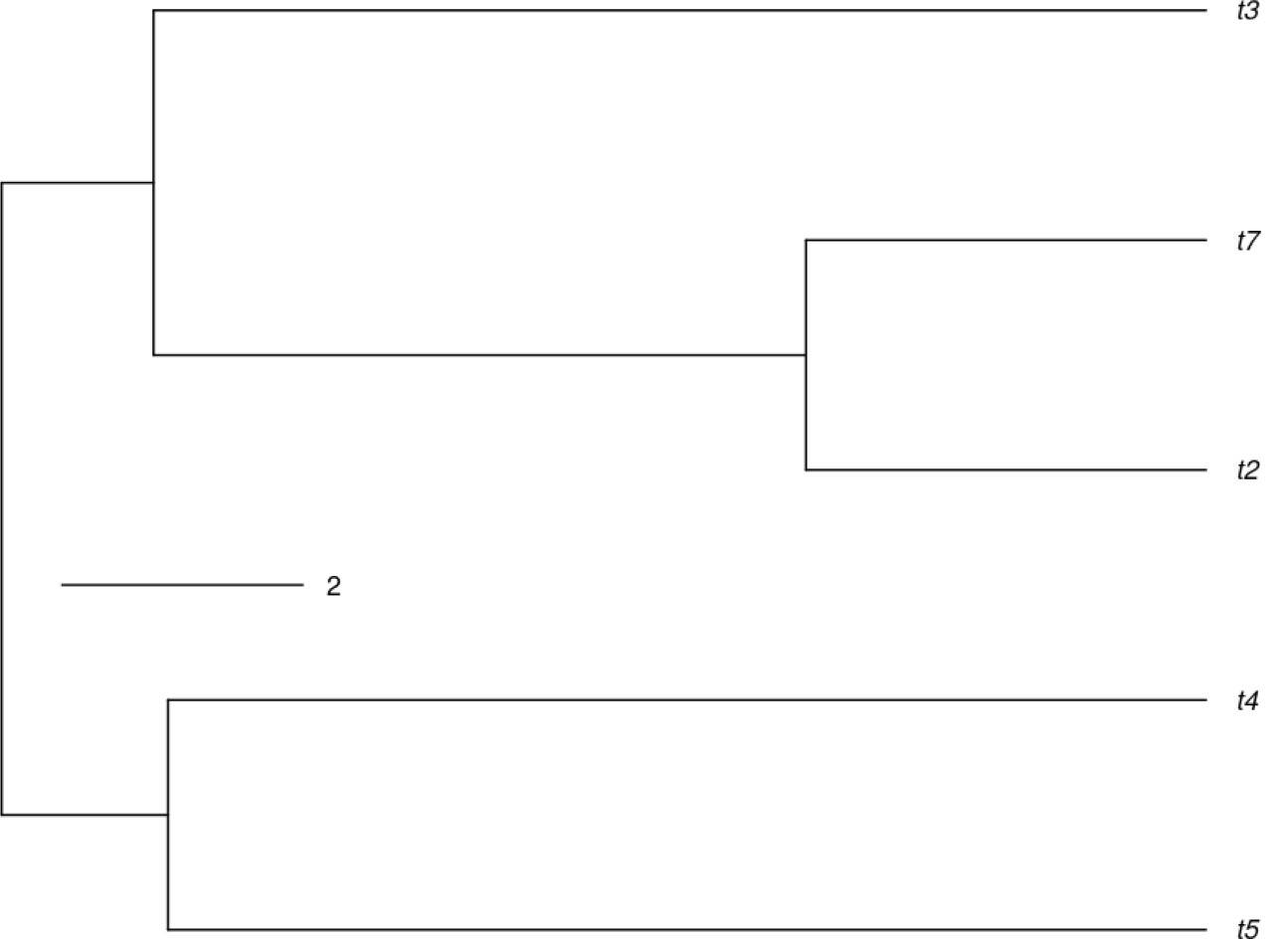
The example tree resulting from a diversity-dependent (DD) simulation.

**Figure 3:**
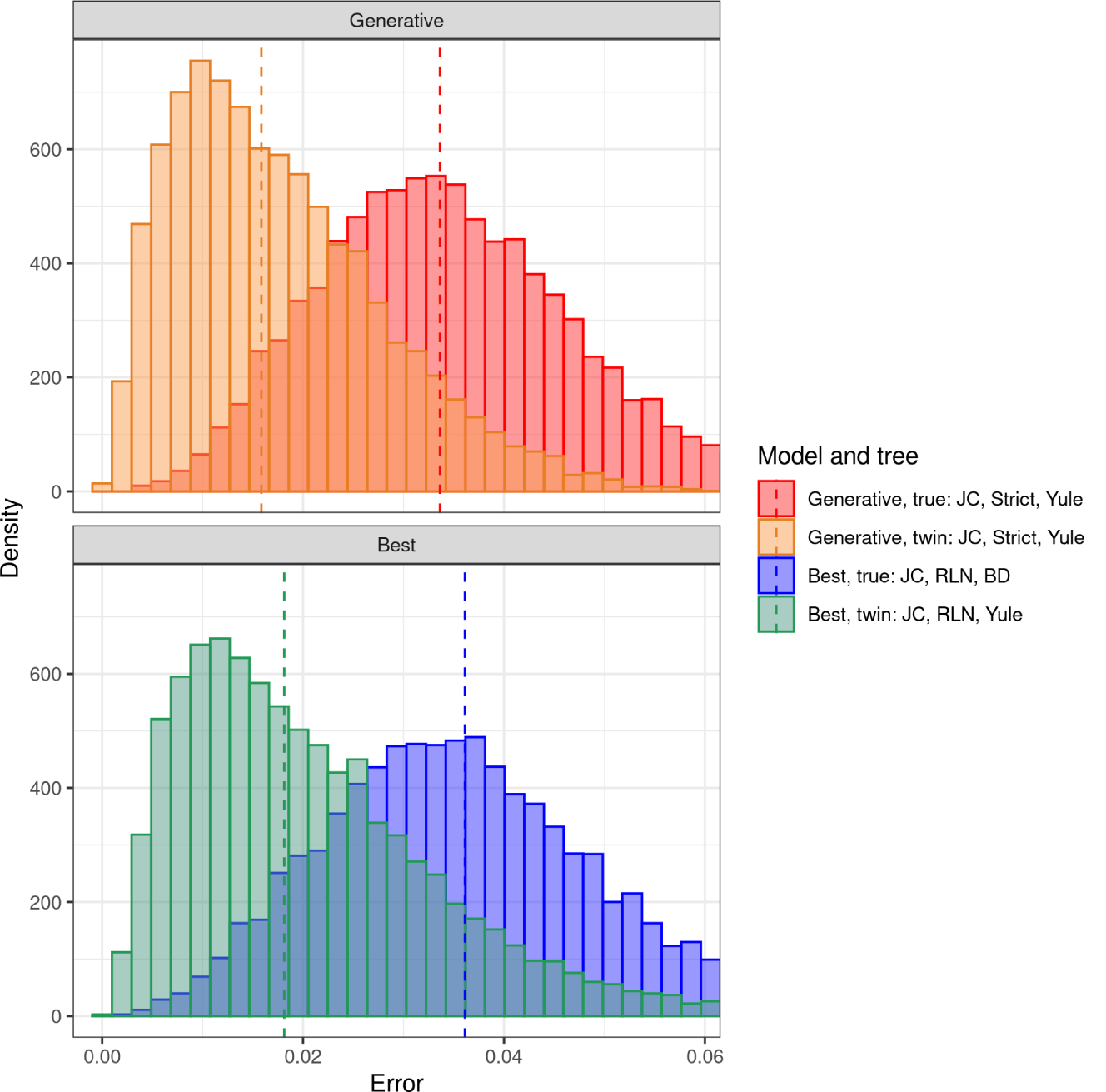
The impact of the tree prior for the example tree in Figure 2. The alignment for this true tree was generated using a JC substitution model and strict clock model. For inferring the tree from this alignment in the ‘generative’ scenario, the same substitution and clock models were used, and a Yule tree prior (this is the assumed generative model, because the real generative model is assumed to be unknown). For the twin tree, the same inference models were used. In the ‘best’ scenario, for the true tree, the best-fitting candidate models were JC substitution model, RLN clock model and BD tree prior, while for the twin tree, the best-fitting candidate models were JC substitution model, RLN clock model and Yule tree prior. The twin distributions show the baseline inference error. Vertical dashed lines show the median error value per distribution.

The third step infers the posterior distributions, using the simulated alignment (Fig. 1, 2a → 4a), and the inference models that were selected in the previous step (3a). For each selected experiment a posterior distribution is inferred, using the `babette` [Bilderbeek & Etienne 2018] R package which makes use of BEAST2.

The fourth step quantifies the newimpact of choosing standard models for inference, i.e. the inference error made. First the burn-in fraction is removed, i.e. the first phase of the Markov chain Monte Carlo (MCMC) run, which samples an unrepresentative part of parameter and tree space. From the remaining posterior, `pirouette` creates an error distribution, by measuring the difference between the true tree and each of the posterior trees (Fig. 1, 4a → 5a). The user can specify a function to quantify the differences between the true and posterior trees.

### 2.2 Controls

`pirouette` allows for two types of control measurements. The first type of control is called ‘twinning’, which results in an error distribution that is the baseline error of the inference pipeline (see supplementary materials, subsection 5.5 for more details). This the error that arises when the models used in inference are identical to the ones used in generating the alignments. The second type of control is the use of candidate models, which result in an error distribution for a generative model that is determined to be the best fit to the tree (see supplementary materials, section 5.6 for more details). The underlying idea is that using a substitution model in inference than used in generating the alignment may partly compensate for choosing a standard tree model instead of the generative tree model as tree prior in inference. Additionally, multiple `pirouette` runs are needed to reduce the influence of stochasticity (see supplementary materials, section 5.7 for more details).

## 3 Usage

We show the usage of `pirouette` on a tree generated by the non-standard diversity-dependent (DD) tree model [Etienne & Haegeman 2020; Etienne *et al*. 2012], which is a BD model with a speciation rate that depends on the number of species.

The code to reproduce our results can be found at https://github.com/richelbilderbeek/pirouette_example_30 and a simplified version is shown here for convenience:

~~~
library (pirouette)
# Create a DD phylogeny with 5 taxa and a crown age of 10
phylogeny <-create_ exemplary _ dd_ tree ()
# Use standard pirouette setup. This creates a list object
   with all settings for generating the alignment, the
   inference using BEAST2, the twinning parameters to
   generate the twin tree and infer it using BEAST2, and the
   error measure
pir_ params <-create_ std _ pir_ params ()
# Do the runs
pir_ out <-pir_ run (
  phylogeny = phylogeny ,
  pir_ params = pir_ params
)
# Plot
pir_ plot(pir_ out)
~~~

The DD tree generated by this code is shown in Figure 2.

The error distribution shown in Figure 3 is produced, which uses the nLTT statistic [Janzen *et al*. 2015] to compare phylogenies (see section 5.8 for details regarding the nLTT statistic and its caveats).

In the upper panel of Figure 3, we can see that the error distributions of the (assumed) generative model (i.e. the known generative substitution and clock models, and the tree model that is assumed in inference of the true tree, and the tree model that is used for generating and inferring the twin tree) differ substantially between the true and twin tree. This difference shows the extent of the mismatch between the true tree model (which is DD) and the (Yule) tree prior used in inference. Because these distributions are distinctively different, the inference error made when using an incorrect tree prior on a DD tree is quite profound.

Comparing the upper and lower panel of Figure 3, we can see that the best candidate model is slightly worse at inferring the true tree, than the (assumed) generative model, indicating that the generative inference model we selected is a good choice.

The candidate model that had highest evidence given the simulated alignment, was JC, RLN and BD (see Table 1 for the meaning of these abbreviations). The RLN clock model is a surprising result: it assumes nucleotide substitutions occur at different rates between the taxa. The JC nucleotide substitution model matches the model used to simulate the alignment. The BD model is perhaps somewhat surprising for the true tree, because the other alternative standard tree prior, Yule, is probably closest to the true DD model because it shows no pull-of-the-present (but also no slowdown).

## 4 Discussion

We showed how to use `pirouette` to quantify the impact of a tree prior in Bayesian phylogenetics, assuming - for illustrative purposes - the simplest standard substitution, clock and tree models, but also the models that would be selected among many different standard tree priors according to the highest marginal likelihood, as this would be a likely strategy for an empiricist. We recommend exploring different candidate models, but note that this is computationally highly demanding, particularly for large trees.

Figure 3 illustrates the primary result of our pipeline: it shows the error distributions for the true tree and the twin tree when either the generative model (for substitution and clock models these are known, for the tree model it must be assumed for the true tree and it is known for the twin tree) or the best-fitting set candidate model (i.e. combination of tree model, substitution model and clock model) is used in inference. The clear difference between the error distributions for the true tree and the twin tree suggests that the choice of tree prior matters. We note, however, that only one tree from a novel tree model is not enough to determine the impact of using an incorrect tree prior. Instead, a distribution of multiple trees, generated by the novel tree model, should be used. In the supplementary material we have provided some examples.

Like most phylogenetic experiments, the setup of `pirouette` involves many choices. A prime example is the length of the simulated DNA sequence. One expects that the inference error decreases for longer DNA sequences. We investigated this superficially and confirmed this prediction (see the supplementary material). However, we note that for longer DNA sequences, the assumption of the same substitution rates across the entire sequence may become less realistic (different genes may experience different substitution rates) and hence longer sequences may require more parameters. Hence, simply getting longer sequences will not always lead to a drastic reduction of the influence of the species tree prior. Fortunately, `pirouette` provides a pipeline that works for all choices.

Interpreting the results of `pirouette` is up to the user; `pirouette` does not answer the question whether the inference error is too large to trust the inferred tree. The user is encouraged to use different statistics to measure the error. The nLTT statistic is a promising starting point, as it can compare any two trees and results in an error distribution of known range, but one may also explore other statistics, for example statistics that depend on the topology of the tree, While `pirouette` allows for this in principle, in our example we used a diversification model (DD) that only deviates from the Yule and BD models in the temporal branching pattern, not in the topology. For models that make different predictions on topology, the twinning process should be modified. As noted in the introduction, Duchêne and colleagues [Duchene *et al*. 2018] also developed a method to assess the adequacy of a tree model on empirical trees. They simulated trees from the posterior distribution of the parameters and then compared this to the originally inferred tree using tree statistics, to determine whether the assumed tree model in inference indeed generates the tree as inferred. This is useful if these trees match, but when they do not, this does not mean that the inferred tree is incorrect; if sufficient data is available the species tree prior may not be important, and hence inference may be adequate even though the assumed species tree prior is not. In short, the approach is applied to empirical trees and compares the posterior and prior distribution of trees (with the latter generated with the posterior parameters!). By contrast, `pirouette` aims to expose when assuming standard priors for the species tree are a mis- or underparameterization. Hence, our approach applies to simulated trees and compares the posterior distributions of trees generated with a standard and non-standard model, but inferred with a standard one. The two methods therefore complement one another. Furthermore, we note that the `pirouette` pipeline is not restricted to exploring the effects of a new species tree model. The pipeline can also be used to explore the effects of non-standard clock or site models, such as relaxed clock models with a non-standard distribution, correlated substitutions on sister lineages, or elevated substitutions rates during speciation events. It is, however, beyond the scope of this paper to discuss all these options in more detail.

In conclusion, `pirouette` can show the errors in phylogenetic reconstruction expected when the model assumed in inference is different from the actual generative model. The user can then judge whether or not this new model should be implemented in a Bayesian phylogenetic tool.

