## Supplementary materials for "Quantifying the impact of an inference model in Bayesian phylogenetics"

### 5 Supplementary material

This supplementary material contains additional facets of `pirouette`, such as the installation of the package, an overview of pirouette's main functions and a guide for users, based on multiple experiments that are shown here as well.

For these experiments, we limited the number of replicates by time, aiming at a duration of 24 hours per setting, when run on the Peregrine computer cluster of the University of Groningen. Due to this, for example, a run of 40 taxa only has few replicates, because one run takes 4 hours. For all experiments, the intermediate results can all be downloaded from their respective websites, which is approximately 5 gigabyte in total.

All the figures shown in this section are shown without any aesthetical modifications, with the exception that the arrangement of the sub-figures in subsection 5.10, where we aligned parts of the figure by hand.

Here is an overview of the various sections:

- subsection 5.1: guidelines for users
- subsection 5.2: installation
- subsection 5.3: resources, such as website, tutorials, packages used, bug reporting and contributing
- subsection 5.4: citation of pirouette
- subsection 5.5: the twinning process
- subsection 5.6: candidate models for the inference
- subsection 5.7: the effects of stochasticity
- subsection 5.8: the nLTT statistic
- subsection 5.9: main functions

- subsection 5.10: code, extra figures and diagnostics regarding the main example.
- subsection 5.11: the result of using multiple trees, as generated by the same stochastic process as the main example
- subsection 5.12: the effect of the number of taxa
- subsection 5.13: the effect of the DNA alignment sequence length
- subsection 5.14 shows the effect when performing inference in the simplest use case
- subsection 5.15 shows the effect when performing inference with an under-parameterization
- subsection 5.17 shows the effect when the twin alignment is allowed to have a different number of substitutions
- subsection 5.18 shows the effect of different mutation rates
- subsection 6: Acknowledgments
- subsection 7: Data accessibility
- subsection 8: Author contributions

#### 5.1 Guidelines for users

From the experiments shown below, we composed some rough guidelines. These guidelines should be treated as preliminary results, as the total runtime of these experiments is 'only' 19 days.

- The use of 20 replicates results in decent plots.
- The use of more taxa increases the inference error

- The use of longer DNA sequences decreases the inference error.
- When we do not impose the same number of substitutions between true and twin alignment, we observe a difference in the error distributions with respect to the standard case (presented in the main text) where they are forced to have the same number of substitutions.
- Using a mutation rate less than 1.0 / crown age, decreases the inference error. We predict this will increase the error in the parameter estimation.

#### 5.2 Installation

`pirouette` will be made available on CRAN from which it can then be easily installed:

```
install.packages("pirouette")
```

Until it is on CRAN, and for the most up-to-date version, one can download and install the package from `pirouette`'s GitHub repository. We first need the `mcbette` and `nodeSub` packages:

```
remotes::install_github(
  "richelbilderbeek/mcbette"
)
remotes::install_github(
  "thijsjanzen/nodeSub"
)
```

Now we can install `pirouette`:

```
remotes::install_github(
  "richelbilderbeek/pirouette"
)
```

which also installs its dependencies from CRAN.

To start using `pirouette`, load its functions in the global namespace first:

```
library(pirouette)
```

Because `pirouette` calls BEAST2, BEAST2 must be installed. This can be done from within R, using:

```
beastier::install_beast2()
```

For the option to select the best candidate model, `pirouette` needs the ”NS” BEAST2 package [Russel *et al.* 2019]. It can be installed from within R, using:

```
mauricer::install_beast2_pkg("NS")
```

#### 5.3 Resources

`pirouette` is free, libre and open source software available at

`http://github.com/richelbilderbeek/pirouette`,

licensed under the GNU General Public License version 3. `pirouette` depends on multiple packages, which are: `ape` [Paradis *et al.* 2004], `assertive` [Cotton 2016], `babette` [Bilderbeek & Etienne 2018], `DDD` [Etienne & Haegeman 2020], `devtools` [Wickham & Chang 2016], `dplyr` [Wickham *et al.* 2019], `ggplot2` [Wickham 2009], `knitr` [Xie 2017], `lintr` [Hester 2016], `magrittr` [Bache & Wickham 2014], `mcbette` [Bilderbeek 2019], `nLTT` [Janzen 2019], `phangorn` [Schliep 2011], `phytools` [Revell 2012], `plyr` [Wickham 2011a], `rappdirs` [Ratnakumar *et al.* 2016], `rmarkdown` [Allaire *et al.* 2017], `Rmpfr` [Maechler 2019], `stringr` [Wickham 2017], `TESS` [Höhna *et al.* 2016b; Höhna 2013], `testit` [Xie 2014], `testthat` [Wickham 2011b] and `tidyr` [Wickham & Henry 2019].

`pirouette`'s development takes place on GitHub,

`https://github.com/richelbilderbeek/pirouette`,

which allows submitting bug reports, requesting features, and adding code. To improve quality, `pirouette` uses a continuous integration service, has a code coverage of above 95% and enforces the most commonly used R style guide [Wickham 2015].

`pirouette`'s is extensively documented on its website, its documentation and its vignettes. The `pirouette` website is a good starting point to learn how to use `pirouette`, as it links to tutorials and videos. The `pirouette` package documentation describes all functions and liberally links to related functions. All exported functions show a minimal example as part of their documentation. The `pirouette` vignette demonstrates extensively how to use `pirouette` in a more informally written way.

The code used in this article and more examples that are periodically tested, can be found at

`https://github.com/richelbilderbeek/pirouette_examples`.

#### 5.4 Citation of pirouette

To cite `pirouette` this article from within R, use:

```
> citation("pirouette")
```

#### 5.5 The twinning process

`pirouette` allows to perform a control measurement, by use of a process we call twinning. This control results in an error distribution that is the baseline error of the pipeline. The difference between the 'true' and 'twin' error distributions is caused only by the mismatch between the true tree model and the tree prior used in the actual inference.

The twinning process, $T$, encompasses two steps: $T_1$, that generates a 'twin tree' (Fig. 1, 1b) and $T_2$, which generates a 'twin alignment' (Fig. 1, 2b). Both twin tree and alignment will be analyzed in the same way as the true tree and alignment.

We define a phylogeny $\tau$ as the combination of branching times $\vec{t}$ and topology $\psi$, and denote as $\tau_G$ the phylogeny produced by a (possibly non-standard) generative diversification model, having branching times $\vec{t}_G$ and topology $\psi_G$.

The first step ($T_1$) of the twinning process creates a tree $\tau_T$ with branching times $\vec{t}_T$ while preserving the original topology $\psi_G$:

$$\tau_G = (\vec{t}_G, \psi_G) \xrightarrow{T_1} \tau_T = (\vec{t}_T, \psi_G) \quad (1)$$

We chose to preserve the original topology to increase the similarity between the twin to the original tree. This works well in the cases of BD or DD models we consider in our example, because all these models make the same assumption about topology (all topologies are equally likely). However, this might not be suitable for new models that assign different probabilities to trees with the same branching times but different topologies. The default option for the twin diversification model $p_T$ is the standard BD model. `pirouette` has a built-in function to use a Yule model as well. Additionally, a user can specify a function to generate a twin tree from any speciation model, such as, for example, a coalescent model.

It is then possible to use the likelihood function $L_T$ for this diversification model to find the parameters $\theta_T^*$ (e.g. speciation and extinction rates, in case of a BD model) that maximize this likelihood applied to the true tree, conditioned on its number of tips $n_G$:

$$\max[L_T(\theta_T | \tau_G, n_G)] \rightarrow \theta_T^*. \quad (2)$$

We use $\theta_T^*$ to simulate a number $n_T = n_G$ of branching times $\vec{t}_T$ for the twin tree $\tau_T$, under the process $p_T$, while preserving the topology. We simulate the new branching times using the TESS package [Höhna *et al.* 2016b]. For simplicity, when simulating phylogenies we assumed a sampling fraction of 100%. A different choice might have an effect on model performance.

The second step ($T_2$) of the twinning process simulates the twin alignment with the same clock model, site model and mutation rate used to simulate the alignment on the true. The twin alignment can be simulated in any user-defined way. `pirouette` provides the option simulate it with the same mutation rate as the true alignment. By default, however, not only the same mutation rate is used, but also the total number of substitutions matches the true alignment. The total number of substitutions is defined as the number of different nucleotides between the (known) root sequence compared to the sequences at the tips.

#### 5.6 Candidate models

The user has to specify exactly one standard inference model, but may be unsure which one to pick. To account for this, the user can specify a set of candidate inference models. Each of these candidate inference models is run in an initial, relatively short, analysis; the candidate model with the highest evidence (i.e., marginal likelihood) will then be used in another, longer, inference run, resulting in another error distribution. The evidence for an inference model is estimated by nested sampling [Russel *et al.* 2019], using the `NS` BEAST2 package.

If twinning is used, a candidate model that has the highest evidence for the twin alignment is also used to create the twin error distribution.

#### 5.7 Stochasticity caused by simulating phylogenies

The goal is to evaluate BEAST2's performance on a non-standard tree model, one must also consider the last source of stochasticity: the different phylogenies a tree model generates. A single phylogeny cannot be considered as fully representative of the model. For this reason multiple phylogenies must be considered (at least 100 independent true and twin trees). If the number of considered phylogenies is high enough, the comparison between the main pipeline's aggregated error distribution and its twin counterpart leads to a fair evaluation of the new tree model with respect to the baseline error.

#### 5.8 The nLTT statistic

The nLTT statistic is the absolute difference between the normalized lineages-through-time plots of two trees. The nLTT statistic is chosen, as it can operate on any two trees (regardless of their crown ages and number of taxa) and its results have a clear range from zero to one. This normalized result makes it possible to compare trees from a distribution of trees from any tree model. The nLTT statistic is not suitable, however, to distinguish between a constant-rate BD model and a family of time-dependent models [Louca & Pennell 2020].

#### 5.9 Main functions

An overview of `pirouette`'s main functions is shown in Table 3. All `pirouette`'s functions are documented, have a useful example and sensible defaults.

| **Name** | **Description** |
|---|---|
| `pir_run` | Run `pirouette` |
| `pir_plot` | Show the `pirouette` results as a plot |
| `create_pir_params` | Create the `pirouette` parameters |
| `create_alignment_params` | Create the alignment parameters |
| `create_twinning_params` | Create the twinning parameters |
| `create_experiment` | Create one experiment |
| `create_error_measure_params` | Create the error measurement parameters |

Table 3: `pirouette`'s main functions and description.

#### 5.10 Main example

This subsection describes the pipeline of the main example and its diagnostics in more detail.

The pipeline starts at the top-left panel of figure 1 (which is identical to figure 2), which is the 'true tree'. The 'true tree' is generated by the diversity-dependent (DD) tree model [Etienne & Haegeman 2020; Etienne *et al.* 2012], which is a BD model with a speciation rate that is dependent on the number of species, with (an arbitrarily chosen) crown age of 10 time units and an expected number of 6 tips for an extinction rate of 0.1. The carrying-capacity is set to 6. The initial speciation rate $\lambda_0$ is chosen such that the expected number of species in a constant-rate BD model would be equal to the number of tips, which amounts to $\lambda_0 = 0.63$. Note that in the main example, a tree was generated with 5 tips, due to stochasticity in the tree generation algorithm.

From this 'true tree', a 'true alignment' is simulated, using the JC nucleotide substitution model and a strict clock model. The resulting alignment is shown at the center-left of figure 1.

From the 'true alignment' the generative inference model is run. Of course, it cannot be the actual (DD) model. Instead, the default BEAST2 inference model is used, which assumes a JC nucleotide substitution model, a strict clock model and a Yule tree model. The resulting posterior trees are shown in the center-left panel of figure 1.

From this 'generative true' posterior (center-left panel in figure 1), the difference between each of its trees is compared to the 'true tree' (top-left panel), using the nLTT statistic, resulting in the error distribution shown in the bottom-left panel of figure 1.

Based on the 'true alignment' (center-left panel), the candidate model with the highest marginal likelihood is determined, from a set of 15 models. The set of models consists of all combinations of all 4 nucleotides substitution models (JC, HKY, TN, GTR), all 2 clock models (strict and relaxed log-normal) and 2 birth-death models (Yule and Birth-Death), except the inference model used as the generative model (JC, strict clock, Yule). The inference model that had the highest evidence (as shown in Table 8) was the inference model with a JC nucleotide substitution model, an RLN clock model and a BD tree model. The resulting posterior trees are shown in the second panel of the third row of posteriors in figure 1.

From this 'best true' posterior, the difference between each of its trees is compared to the 'true tree' (top-left panel), using the nLTT statistic, resulting in the second error distribution in the bottom row of figure 1.

From the 'true tree' (top-left) we generated a BD twin tree (top-right).

From this 'twin tree', a 'twin alignment' was simulated, using the JC nucleotide substitution model and a strict clock model. The resulting alignment is shown in the center-right panel of figure 1.

From the 'twin alignment' the generative inference model is run as well. Also here, the default BEAST2 inference model is used, which assumes a JC nucleotide substitution model, a strict clock model and a Yule tree model. The resulting posterior trees are shown in the third panel of the third row of figure 1. From this 'generative twin' posterior, the difference between each of its trees is compared to the 'twin tree' (top-right panel), using the nLTT statistic, resulting in the error distribution shown in the third panel of the bottom row of figure 1.

Based on the 'twin alignment' (center-right panel), the candidate model with the highest marginal likelihood is determined, from the same set of 15 candidate models. The inference model that had the highest evidence (as shown in Table 9) was the inference model with a JC nucleotide substitution model, an RLN clock model and a Yule tree model (Note that this is different from how the twin tree was generated which was with a BD process and the alignment was simulated with a JC substitution model and strict clock model ). However, the extinction rate used in simulating the twin tree was practically 0, thus resembling a Yule process. From the 'twin alignment' this best candidate inference model is run. The resulting posterior trees are shown in the fourth panel of the third row of posteriors in figure 1.

From this 'best twin' posterior (fourth in third row of figure 1), the difference between each of its trees was compared to the 'twin tree' (top-right panel), using the nLTT statistic, resulting in the fourth error distribution in the bottom row of figure 1.

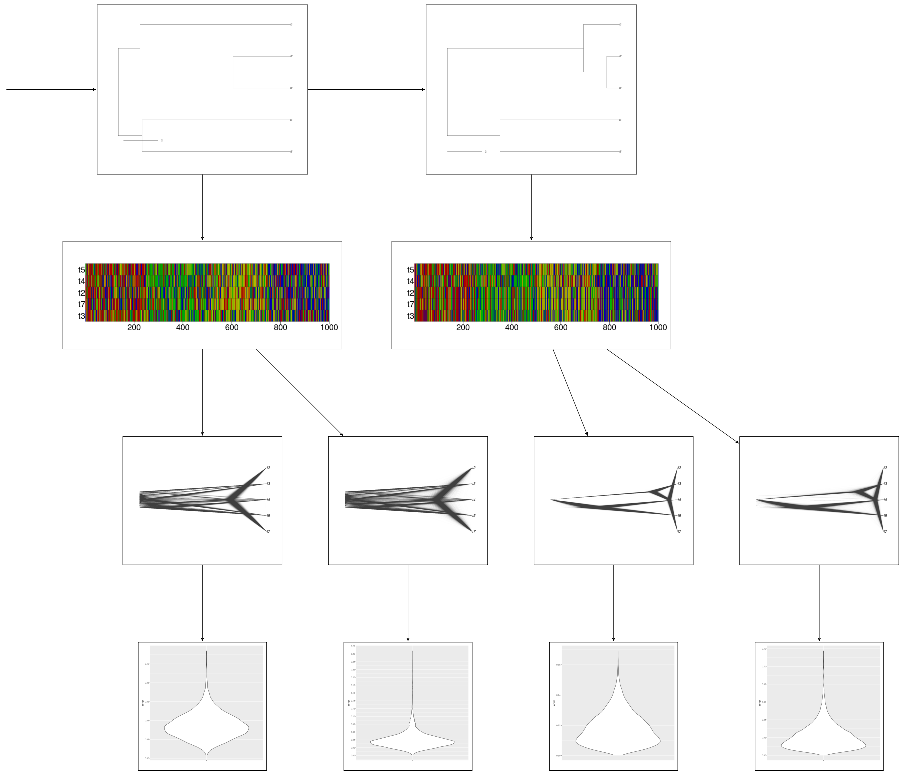


Figure 4: Full pirouette pipeline, including comparison to baseline error. The true tree (top left) is used to simulate an alignment. From this alignment two posterior distributions of trees are created: one using the generative model and another one using the inference model with the highest marginal likelihood. For each distribution of trees, a distribution of errors, measured with the nLTT statistic, between the posterior trees and the main trees is drawn. From the true tree also a twin tree is created (right side of the figure) which follows the same pipeline, leading to two additional error distributions to use as baseline errors.

To assess if the results of the inference are meaningful one important parameter is the Effective Sample Size (ESS). This quantity describes how many independent trees are sampled from the posterior distributions. For reliable results it is good practice to have at least $ESS = 200$ (see `https://beast.community/ess_tutorial`).

In the following we present the ESS for the posterior distributions of the 4 cases shown in Fig. 4.

The ESSes of the 'true' pipeline for the generative model are shown in Table 4. From the estimated parameters, one can deduce that the JC nucleotide substitution model was used (no estimated parameter needed), a strict clock model was used (again, no parameter needed to be estimated) and a Yule tree prior is used ('Yule model' and 'birthRate' are estimated). Note that although the actual true tree is created by a DD process, the default and standard Yule tree model is used as the closest standard tree model.

| parameter | ESS |
|---|---|
| posterior | 10001 |
| likelihood | 10001 |
| prior | 9804 |
| treeLikelihood | 10001 |
| TreeHeight | 10001 |
| YuleModel | 9804 |
| birthRate | 9931 |

Table 4: ESSes for generative model

The ESSes of the 'twin' pipeline for the generative model are shown in Table 5. Note that the generative inference model is re-used (which assumes a Yule tree model) in the inference, where the twin tree is actually created using a BD process, which is the default.

| parameter | ESS |
|---|---|
| posterior | 9969 |
| likelihood | 9997 |
| prior | 9955 |
| treeLikelihood | 9997 |
| TreeHeight | 9762 |
| YuleModel | 9955 |
| birthRate | 9844 |

Table 5: ESSes for generative model, twin tree

The ESSes of the 'true' pipeline for the best candidate model are shown in Table 6. From the names of the estimated parameters, it is clear that the best candidate model has a JC nucleotide substitution model (no parameter needed to be estimated) an RLN clock model (which can be inferred from the parameter 'rate.mean') and a BD tree prior ('BirthDeath', 'BDBirthRate' and 'BDDeathRate').

| parameter | ESS |
|---|---|
| posterior | 8320 |
| likelihood | 10001 |
| prior | 2278 |
| treeLikelihood | 10001 |
| TreeHeight | 3239 |
| ucldStdev | 1027 |
| rate.mean | 1853 |
| rate.variance | 543 |
| rate.coefficientOfVariation | 1215 |
| BirthDeath | 7068 |
| BDBirthRate | 7615 |
| BDDeathRate | 6402 |

Table 6: ESSes for best candidate model

The ESSes of the 'twin' pipeline for the best candidate model are shown in Table 7.

From the names of the estimated parameters, it is clear that the best candidate model for the twin tree is JC nucleotide substitution model (no parameter needed to be estimated), an RLN clock model (which can be inferred from the parameter 'rate.mean') and a Yule model ('YuleModel', 'birthRate'). Note that there is a mismatch between the actual process of how the twin tree and twin alignment are generated, as the twin tree is generated by a BD process, and the alignment is simulated using a JC nucleotide substitution model and a strict clock model. Again we note that the extinction rate used to simulate the twin tree (estimated from the true tree) was practically 0, so the BD process resembled a Yule process.

| parameter | ESS |
|---|---|
| posterior | 9623 |
| likelihood | 10001 |
| prior | 2302 |
| treeLikelihood | 10001 |
| TreeHeight | 4513 |
| ucldStdev | 1414 |
| rate.mean | 2923 |
| rate.variance | 1560 |
| rate.coefficientOfVariation | 1625 |
| YuleModel | 7854 |
| birthRate | 9636 |

Table 7: ESSes for best candidate model, twin tree

The marginal likelihood (or evidence) data for the model comparison performed in the 'true' pipeline is shown in Table 8. The best (that is, the one with the highest model weight) candidate model assumes a JC nucleotide substitution model, an RLN clock and a BD tree model.

| Site model | Clock model | Tree prior | log(evidence) | log(evidence error) | Weight | ESS |
|---|---|---|---|---|---|---|
| GTR | RLN | BD | -6661.105 | 5.895 | 0.000 | 11.422 |
| GTR | RLN | Yule | -6650.211 | 4.669 | 0.000 | 8.311 |
| GTR | Strict | BD | -6656.726 | 5.304 | 0.000 | 11.711 |
| GTR | Strict | Yule | -6656.272 | 5.567 | 0.000 | 7.415 |
| HKY | RLN | BD | -6640.067 | 4.187 | 0.001 | 5.982 |
| HKY | RLN | Yule | -6642.854 | 4.641 | 0.000 | 5.865 |
| HKY | Strict | BD | -6661.308 | 5.857 | 0.000 | 9.510 |
| HKY | Strict | Yule | -6646.973 | 5.396 | 0.000 | 5.643 |
| JC | RLN | BD | -6633.353 | 2.969 | 0.945 | 5.770 |
| JC | RLN | Yule | -6636.447 | 3.363 | 0.043 | 6.941 |
| JC | Strict | BD | -6639.650 | 4.161 | 0.002 | 5.476 |
| TN | RLN | BD | -6640.669 | 3.778 | 0.001 | 6.728 |
| TN | RLN | Yule | -6644.276 | 4.797 | 0.000 | 7.155 |
| TN | Strict | BD | -6638.117 | 3.277 | 0.008 | 5.949 |
| TN | Strict | Yule | -6644.336 | 3.838 | 0.000 | 7.806 |

Table 8: Evidences for the true phylogeny

The marginal likelihood (or evidence) data for model comparison performed in the 'twin' pipeline is shown in Table 9. The best (that is, the one with the highest model weight) candidate model assumes a JC nucleotide substitution model, an RLN clock and a Yule tree model. Note that there is a mismatch between the actual process of how the twin tree and twin alignment are generated, as the twin tree is generated by a BD process, and the alignment is simulated using a JC nucleotide substitution model and a strict clock model. The extinction rate in simulating the BD process (estimated from the true tree) was, however, practically 0, so the BD process resembled a Yule process.

| Site model | Clock model | Tree prior | log(evidence) | log(evidence error) | Weight | ESS |
|---|---|---|---|---|---|---|
| GTR | RLN | BD | -5571.348 | 6.569 | 0.000 | 14.874 |
| GTR | RLN | Yule | -5557.738 | 5.566 | 0.000 | 5.352 |
| GTR | Strict | BD | -5547.850 | 4.386 | 0.000 | 5.312 |
| GTR | Strict | Yule | -5553.065 | 5.664 | 0.000 | 6.393 |
| HKY | RLN | BD | -5548.450 | 3.356 | 0.000 | 12.060 |
| HKY | RLN | Yule | -5544.146 | 3.817 | 0.000 | 4.167 |
| HKY | Strict | BD | -5559.866 | 6.078 | 0.000 | 6.766 |
| HKY | Strict | Yule | -5565.050 | 6.695 | 0.000 | 5.779 |
| JC | RLN | BD | -5536.454 | 2.784 | 0.040 | 8.767 |
| JC | RLN | Yule | -5533.284 | 2.679 | 0.959 | 6.058 |
| JC | Strict | BD | -5541.446 | 3.685 | 0.000 | 4.370 |
| TN | RLN | BD | -5557.132 | 4.844 | 0.000 | 9.823 |
| TN | RLN | Yule | -5557.777 | 4.589 | 0.000 | 7.928 |
| TN | Strict | BD | -5545.206 | 4.229 | 0.000 | 9.899 |
| TN | Strict | Yule | -5546.328 | 3.962 | 0.000 | 9.313 |

Table 9: Evidences for twin phylogeny

#### 5.11 Using a distribution of trees

This subsection extends the main example, by using multiple (instead of one) trees. These trees are produced by running a DD tree simulation with the same parameters as the main example.

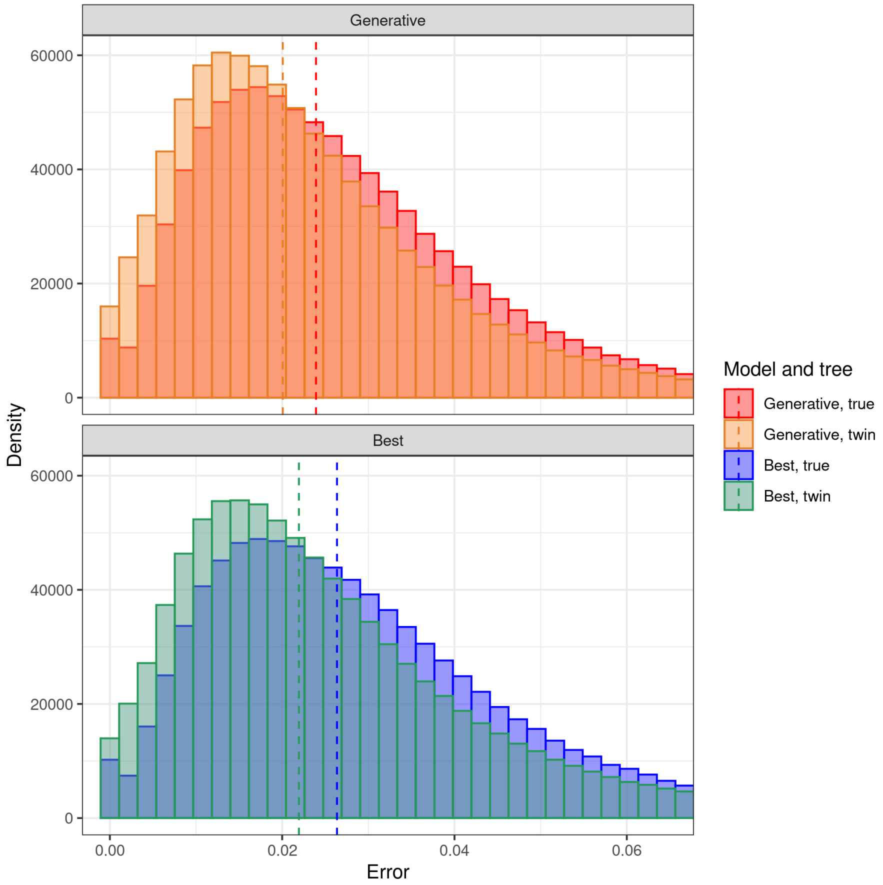


Figure 5: Aggregate error distributions, similar to Fig. 3 for the main example, but now for a collection of 100 replicate trees. For each setting (true generative, true best candidate, twin generative and twin best candidate), the resulting errors from each replicate pipeline have been merged into a single distribution. This took 2.7 days (wall clock time) to compute.

The resulting error distributions are shown in Fig. 5. We present results for cases where (1) the generative model has been used or (2) the model with highest evidence has been selected for the inference. From the plots we can see that in both cases the two distributions (true and twin) are mostly overlapping, but not everywhere. This suggests that the inference models that have been used can to a reasonable extent capture in an accurate way the features of the diversity-dependent tree prior used to simulate the original trees.

The code to reproduce Fig. 5 can be found at

`https://github.com/richelbilderbeek/pirouette_example_28`.

#### 5.12 The effect of the number of taxa

The main example uses 5 taxa. Here we show the same results as the main example, except for a varying number of taxa. We did so, by setting the DD model's carrying capacity to the desired number of taxa.

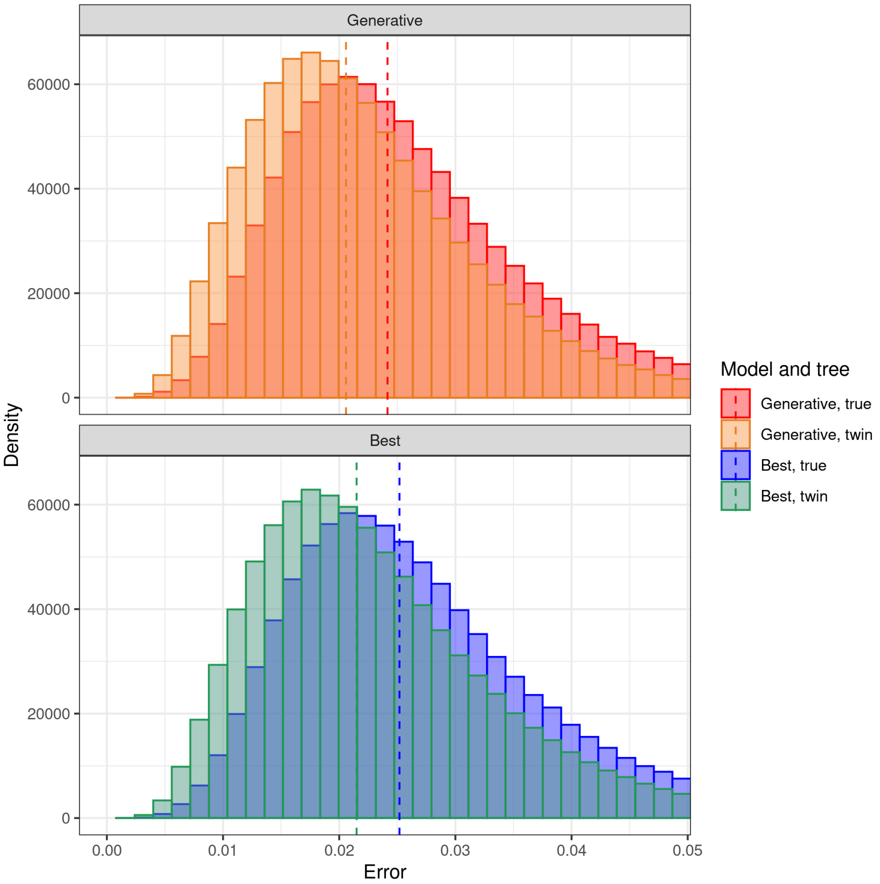


Figure 6: Aggregate error distributions for 100 replicates. Here each true tree has 12 taxa. This took 6.0 days (wall clock time) to compute.

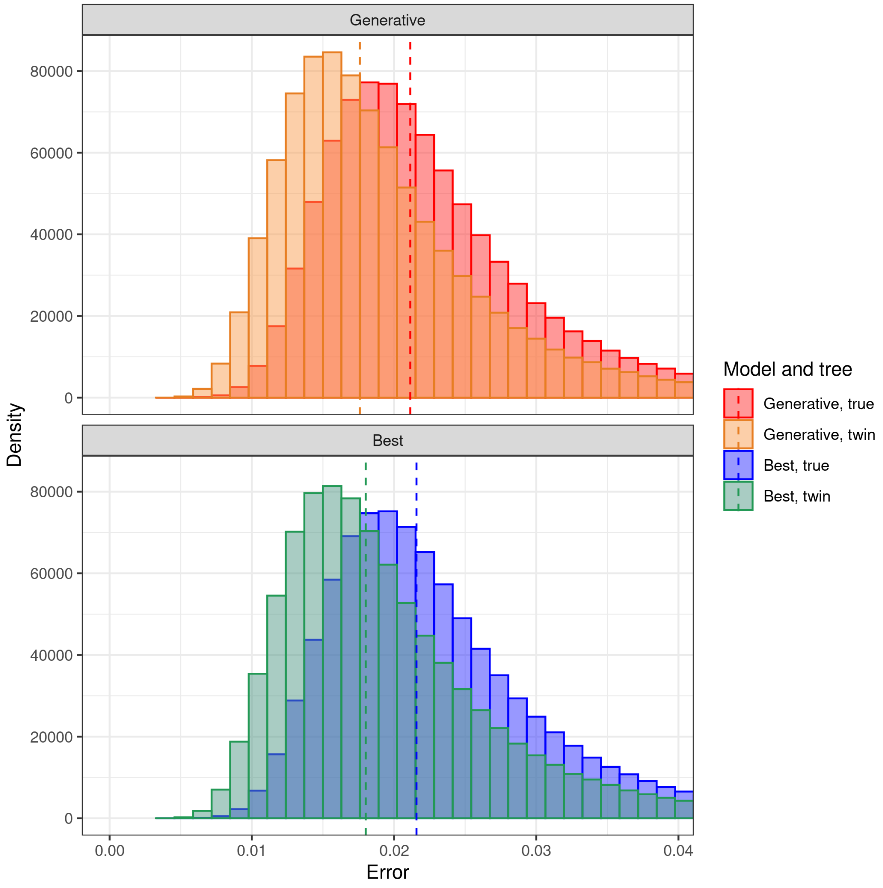


Figure 7: Aggregate error distributions for 100 replicates. Here each true tree has 24 taxa. This took 9.8 days (wall clock time) to compute.

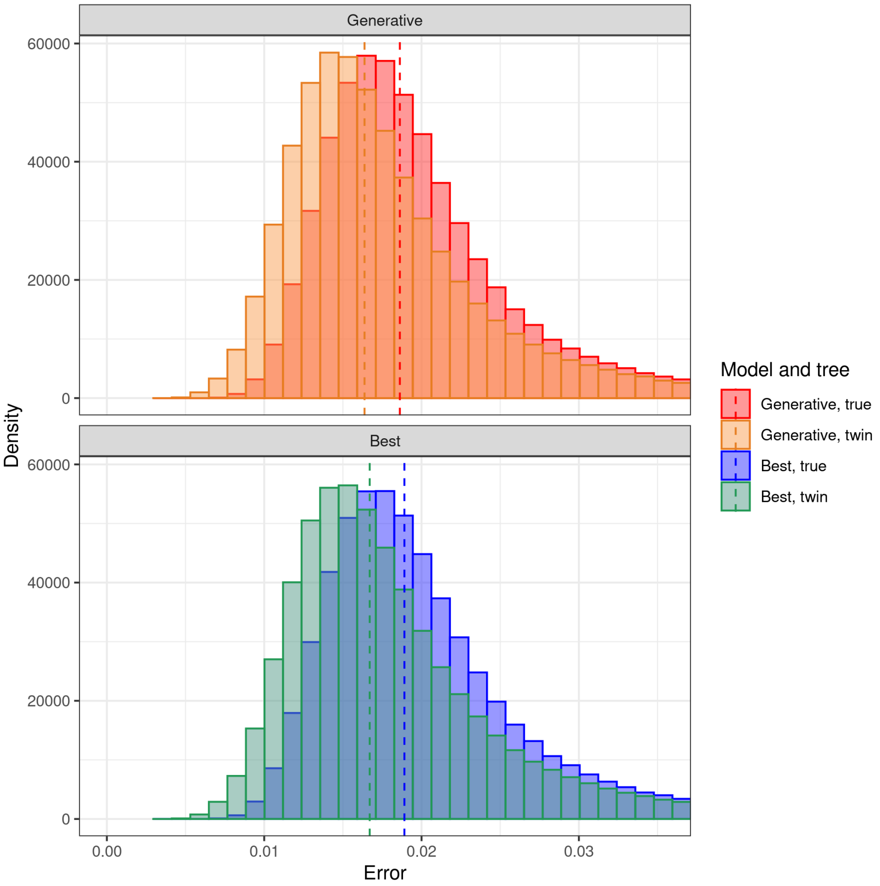


Figure 8: Aggregate error distributions for 65 replicates. Here each true tree has 32 taxa. This took 8.0 days (wall clock time) to compute.

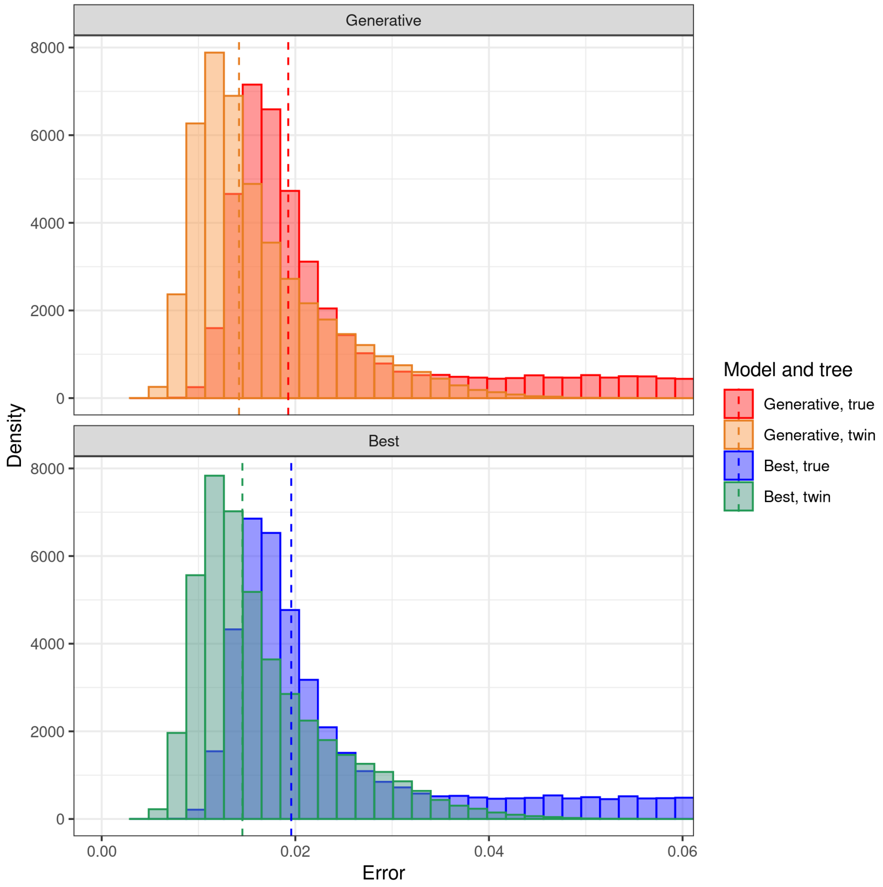


Figure 9: Aggregate error distributions for 5 replicates. Here each true tree has 40 taxa. This took 0.83 days (wall clock time) to compute.

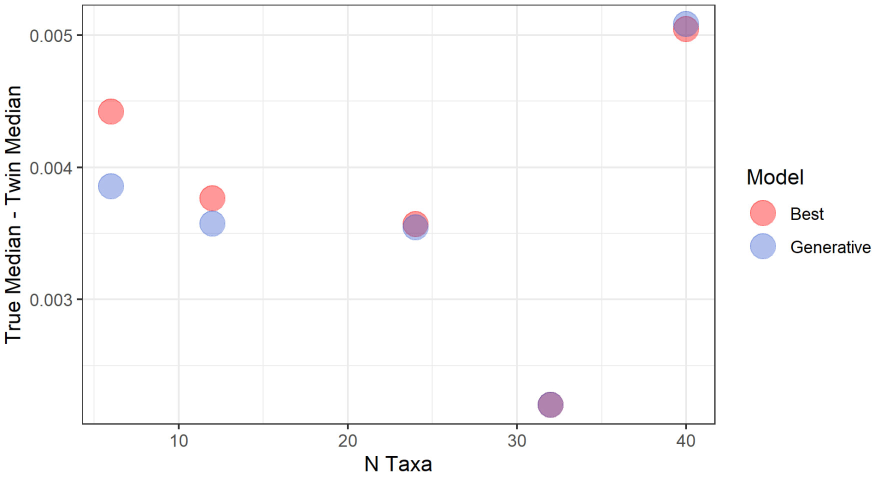


Figure 10: Difference between median true error and median twin error for different number of taxa.

We show in figures 5, 6, 7, 8 and 9 what are the errors obtained when starting from phylogenies with, respectively, 5, 12, 24, 32 and 40 taxa. Again we can see that in each case errors tend to be greater in the true distribution than in the twin distribution, similar to the result of subsection 5.11. Collecting all the data together we can see that errors tend to decrease as the number of taxa in the considered phylogenies increase (see Fig. 10). The data point for 40 taxa not following the trend could be due to the limited amount of simulated trees taken in consideration due to time constraints.

The code to reproduce these figures can be found at `https://github.com/richelbilderbeek/pirouette_example_28` (5 taxa, main example), `https://github.com/richelbilderbeek/pirouette_example_32` (12 taxa), `https://github.com/richelbilderbeek/pirouette_example_33` (24 taxa), `https://github.com/richelbilderbeek/pirouette_example_41` (32 taxa), `https://github.com/richelbilderbeek/pirouette_example_42` (40 taxa).

#### 5.13 The effect of DNA sequence length

The main example uses a DNA alignment length of 1000 nucleotides. Here, we show the same results as the main example, except for a varying DNA alignment sequence length.

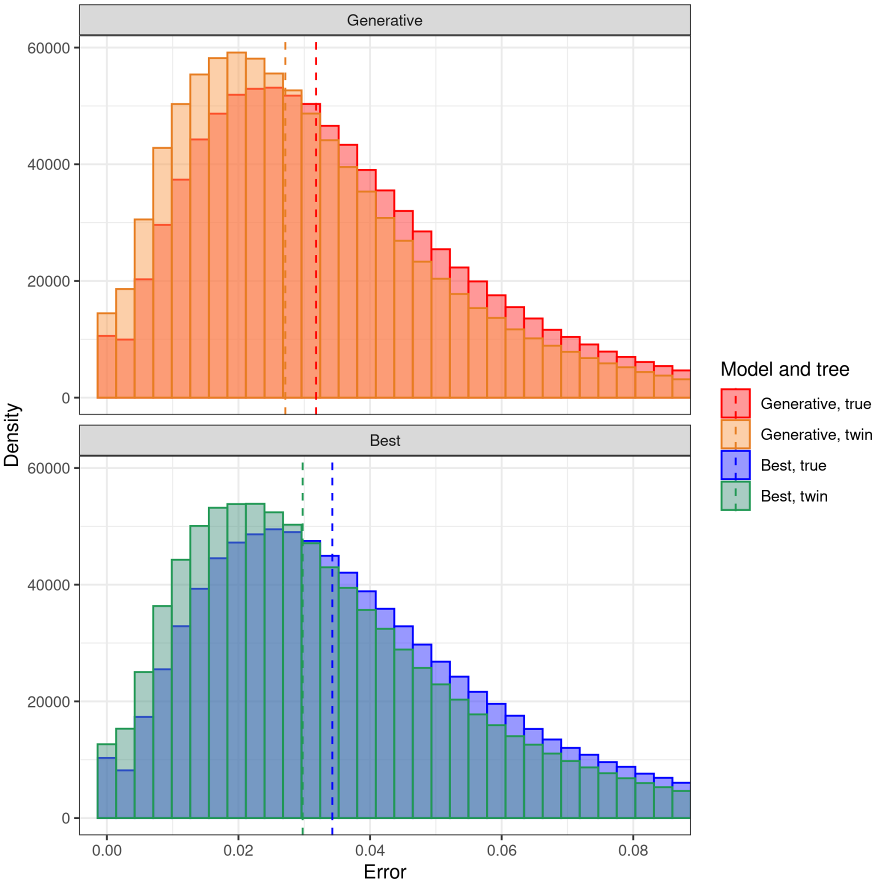


Figure 11: Aggregate error distributions for 100 replicates. Here each each alignment has a sequence length of 500 nucleotides. This took 2.2 days (wall clock time) to compute.

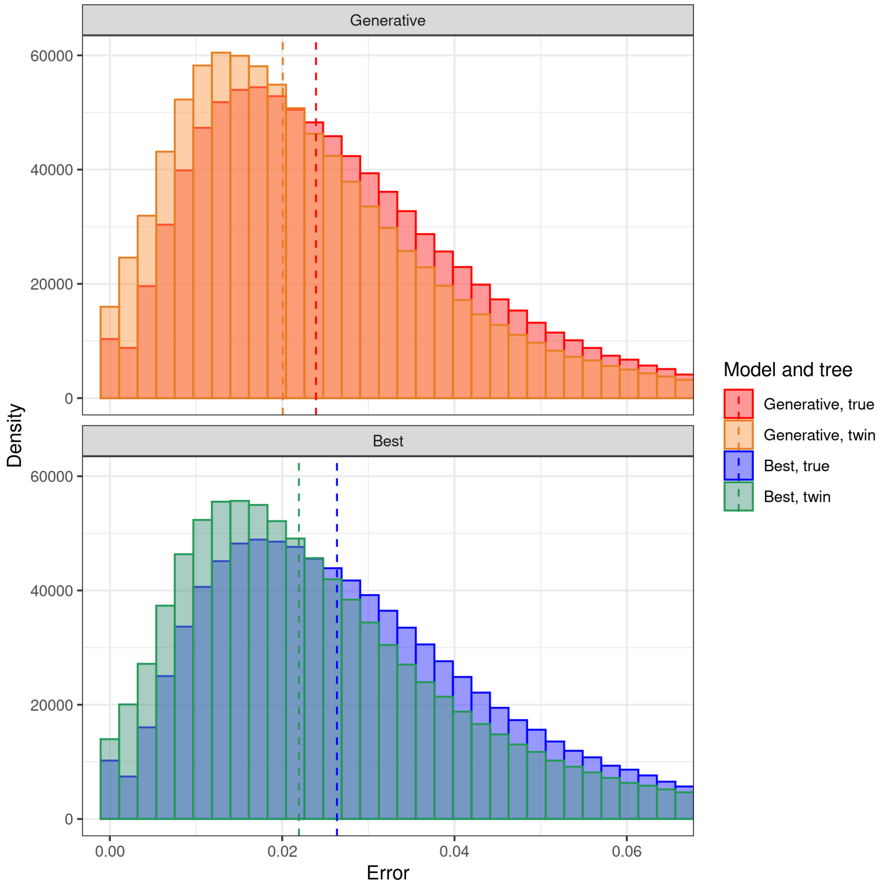


Figure 12: Aggregate error distributions for 100 replicates. Here each each alignment has a sequence length of 1000 nucleotides. This is a replicate of Fig. 5. We put it here to facilitate the comparison with the cases with different number of nucleotides.

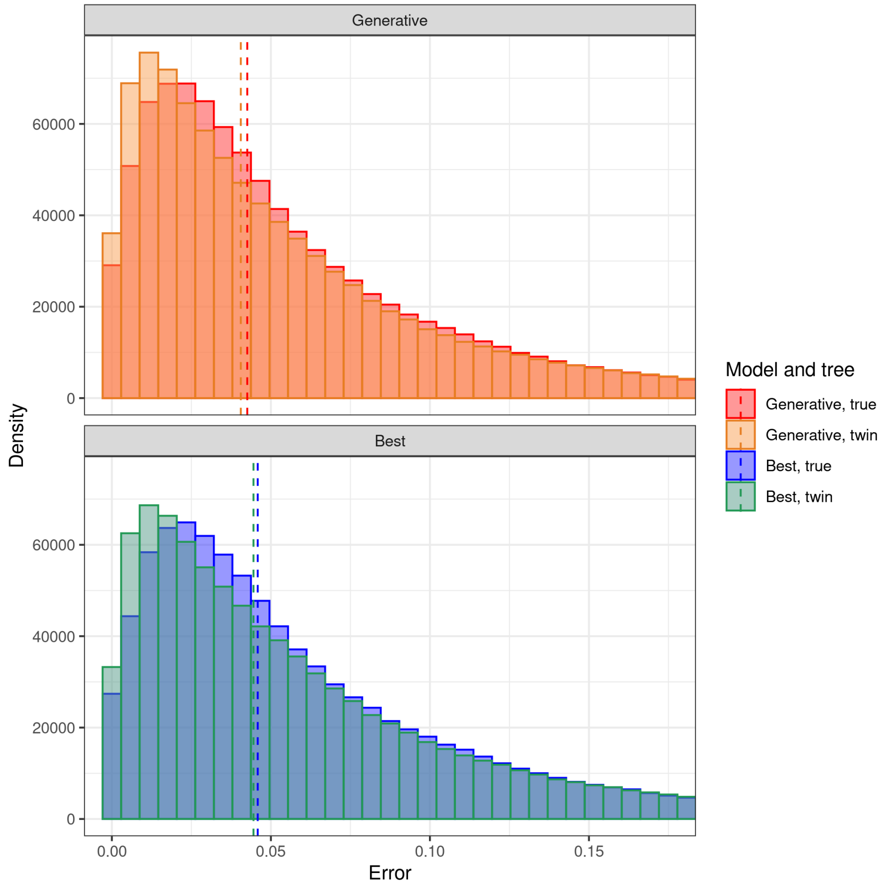


Figure 13: Aggregate error distributions for 100 replicates. Here each each alignment has a sequence length of 2000 nucleotides. This took 4.4 days (wall clock time) to compute.

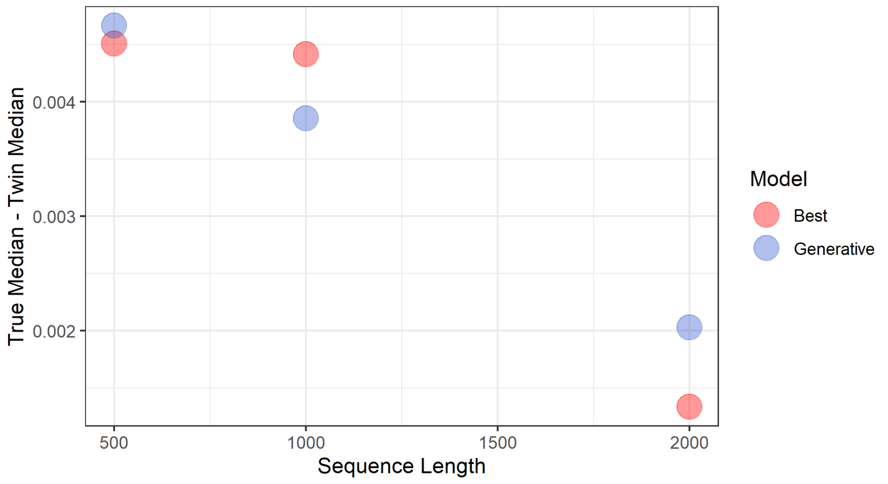


Figure 14: Difference between median true error and median twin error for different sequence lengths.

From figures 11, 12 and 13 we can observe that the discrepancy between the true and twin error distributions tends to become smaller as the number of nucleotides increase (see also Fig. 14). This occurred for both the generative and best candidate cases. This follows the expectation that a prior becomes less important when more information becomes available.

The code to reproduce these figures can be found at `https://github.com/richelbilderbeek/pirouette_example_19` (500 nucleotides), `https://github.com/richelbilderbeek/pirouette_example_28` (1000 nucleotides, main example), and `https://github.com/richelbilderbeek/pirouette_example_34` (2000 nucleotides).

#### 5.14 The effect of assuming a Yule tree prior on a Yule tree

The main example uses a tree generated by a non-standard tree model. Here, we show the same results, with the only difference that the tree used is generated by simplest tree model (the Yule model), which we also assume as the (correct) tree prior.

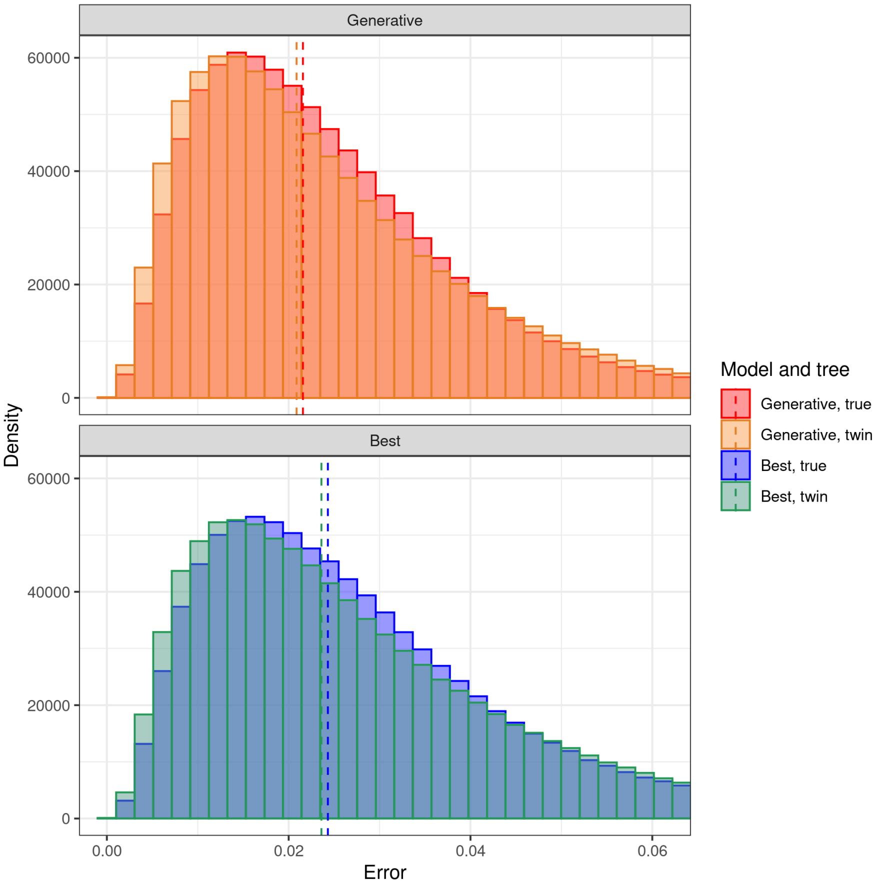


Figure 15: Aggregate error distributions for 100 replicates. Here each true tree is generated by a Yule process. For the inference we used a Yule tree prior. This took 2.9 days (wall clock time) to compute.

This example shows a parameterization at the correct level for the simplest case possible.

As expected the twin and true distributions in Fig. 15 are extremely similar for both the generative and the best candidate case.

The code to reproduce this figure can be found at `https://github.com/richelbilderbeek/pirouette_example_22`.

#### 5.15 The effect of assuming a Yule tree prior on a BD tree

The main example uses a tree generated by a non-standard tree model. Here, we show the same results, with the difference that the tree used is generated by a birth-death (BD) tree model, where we assume it is generated by a Yule (or pure-birth) model. This example thus shows the effect of underparameterization.

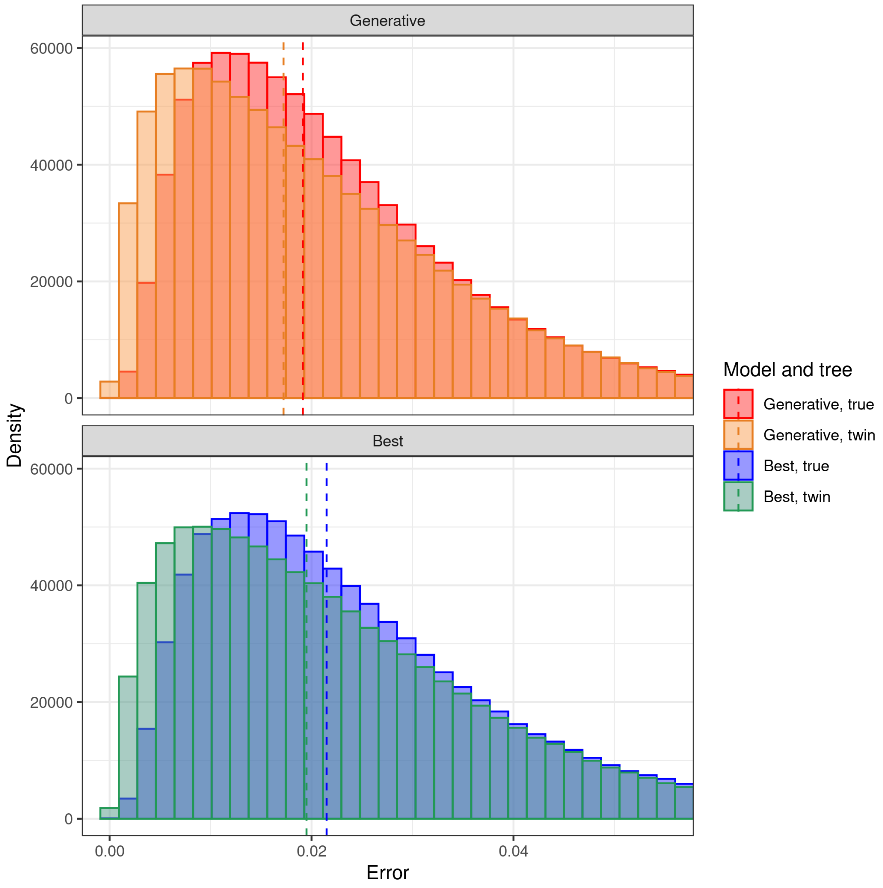


Figure 16: Aggregate error distributions for 100 replicates. Here each true tree is generated by a BD process. For the inference we used instead a Yule tree prior. This took 2.7 days (wall clock time) to compute.

Because the two models are very similar to each other (the BD model can be turned into a Yule model just by setting the extinction parameter to zero [Nee *et al.* 1994]) the median discrepancy is almost negligible. However, with respect to the previous case (subsection 5.14), where a Yule tree prior was used, the distributions here exhibit a greater difference. As we use only extant trees, it is reasonable that the method is slightly weaker in distinguishing between the Yule and BD models. It is unknown what the discriminatory power would be when comparing trees with extinction events.

The code to reproduce this figure can be found at

`https://github.com/richelbilderbeek/pirouette_example_26`.

#### 5.16 The effect of diversity-dependent trees differing in how likely they are under the DD process

Here we show the results of a `pirouette` run on a dataset of multiple DD trees that we selected for having a low, median and high likelihood. In this way, we effectively selected for trees that are rare, uncommon and common respectively.

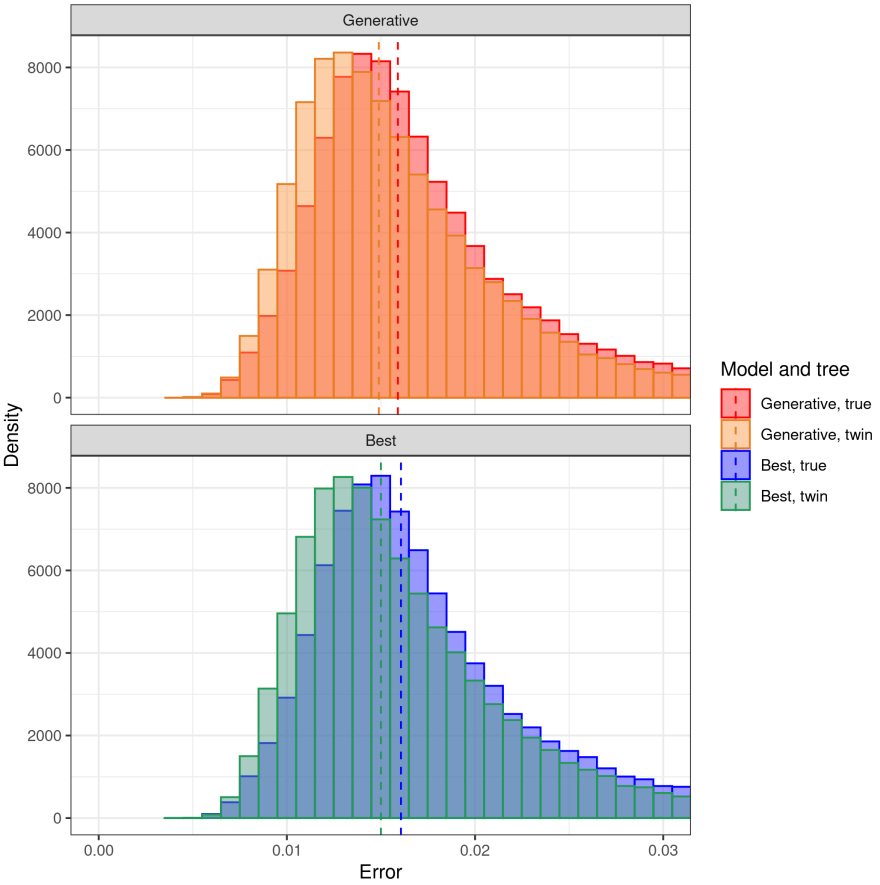


Figure 17: Aggregate error distributions for a distribution of trees, where the true trees are DD with low likelihood.

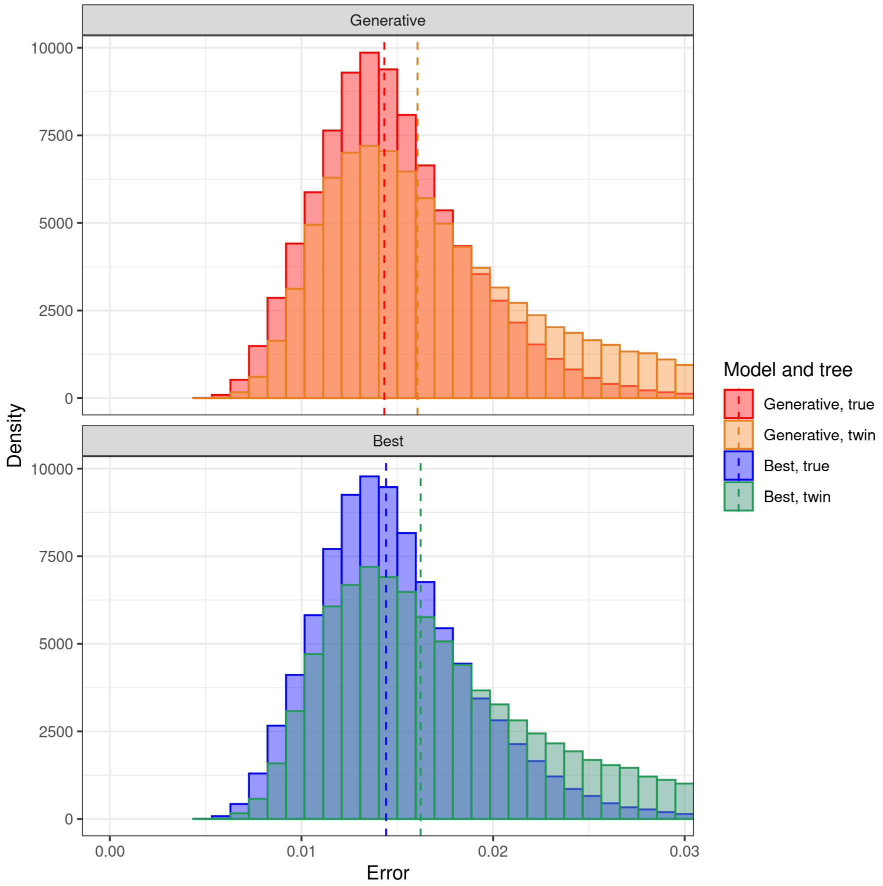


Figure 18: Aggregate error distributions for a distribution of trees, where the true trees are DD with median likelihood.

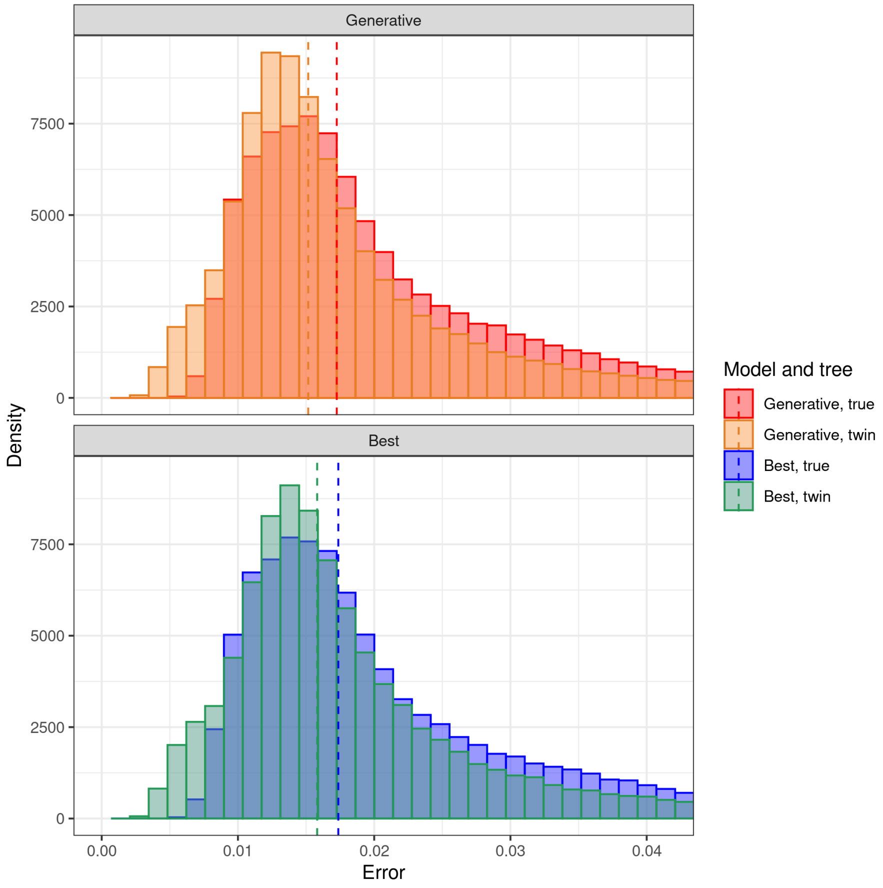


Figure 19: Aggregate error distributions for a distribution of trees, where the true trees are DD with high likelihood.

Here the median errors are similar in the three settings and similar to ones relative to the full dataset of 5.11. We can also notice that in the case of median likelihood, the twin median error appears to be lower than the true mean error. This is usually a sign that the number of replicates (in this case 10) is too low to allow us to draw precise conclusions from this test. We did not explore further in this direction using more simulations because computational times turned to be extremely high. The entire run took 120 hours in total.

The code to reproduce these figure can be found at https://github.com/richelbilderbeek/pirouette_example_23 .

#### 5.17 The effect of equal or equalized mutation rate in the twin alignment

The main example uses a twin alignment that has the same number of substitutions (as measured from the ancestral sequence) as the true alignment. Here, we show the same results, with the difference that the twin alignment uses the same mutation rate, yet is not guaranteed to have the same number of substitutions.

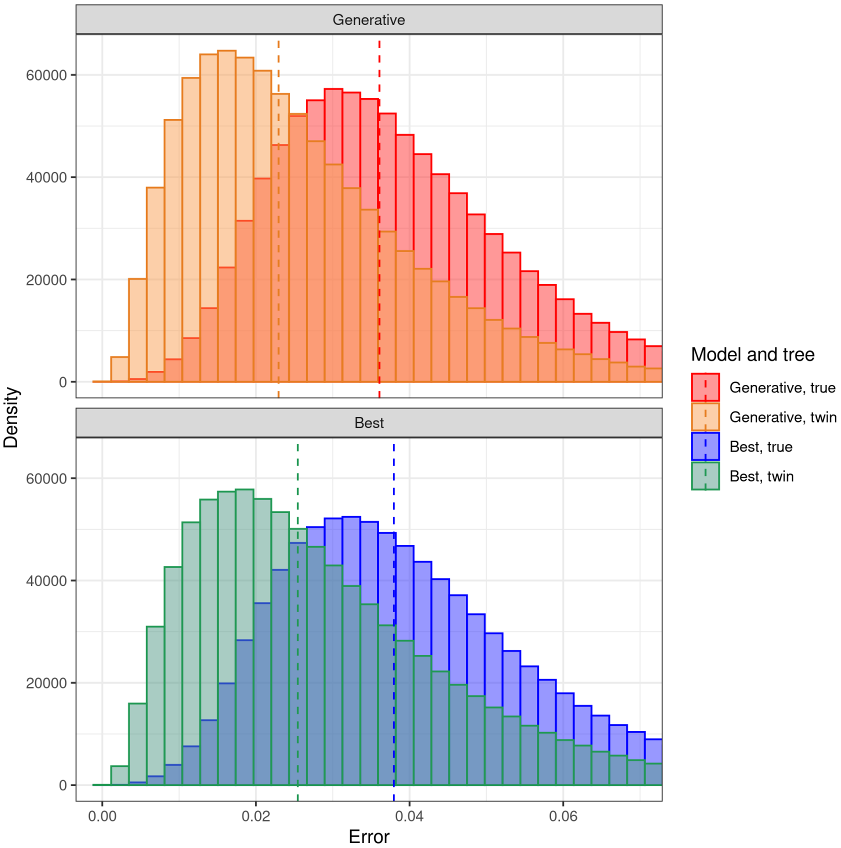


Figure 20: Aggregate error distributions for 100 replicates. similar to Fig. 5, but here the number of substitutions is not imposed to be the same between true and twin alignment. Instead, an equal mutation rate is used. This took 3.3 days (wall clock time) to compute.

Comparing figures 20 and 5 we can see that the discrepancy between true and twin distributions tend to increase. This is probably due to the fact that letting mutation rates induces a difference in the amount of information contained in the alignments and this is reflected in the error distributions.

The code to reproduce this figure can be found at `https://github.com/richelbilderbeek/pirouette_example_18` and `https://github.com/richelbilderbeek/pirouette_example_28`.

##### 5.18 The effect of mutation rate

The main example uses a mutation rate such that all nucleotides, on average, mutate once over the history going from the ancestral sequence at the crown to the alignments at the tips. This value equals '1.0 / crown age'. In this way, the alignment is expected to contain the maximum amount of information.

Here, we show the same results for different mutation rates. The results for the different mutation rates are shown in Figs. 21 (0.25 / crown age), 22 (0.50 / crown age), 23 (0.75 / crown age), 5 (1.00 / crown age), 24 (1.25 / crown age), 25 (1.50 / crown age) and 26 (2.00 / crown age). Fig. 27 summarizes all the other figures showing on the y-axis, for each value of the mutation rate, the difference between the median of the true distribution and the median of the twin distribution. We can observe a general positive trend as the mutation rate increase, even though the value for 1.5 / crown age suggests to take this result with caution. It is possible, however, that a more regular trend could be observed increasing the number of simulations.

The code to reproduce this figure can be found at `https://github.com/richelbilderbeek/pirouette_example_35` (0.25 / crown age), `https://github.com/richelbilderbeek/pirouette_example_36` (0.50 / crown age), `https://github.com/richelbilderbeek/pirouette_example_37` (0.75 / crown age), `https://github.com/richelbilderbeek/pirouette_example_28` (1.00 / crown age, example reported in 5.11, see Fig. 5), `https://github.com/richelbilderbeek/pirouette_example_38` (1.25 / crown age), `https://github.com/richelbilderbeek/pirouette_example_39` (1.50 / crown age), `https://github.com/richelbilderbeek/pirouette_example_40` (2.00 / crown age),

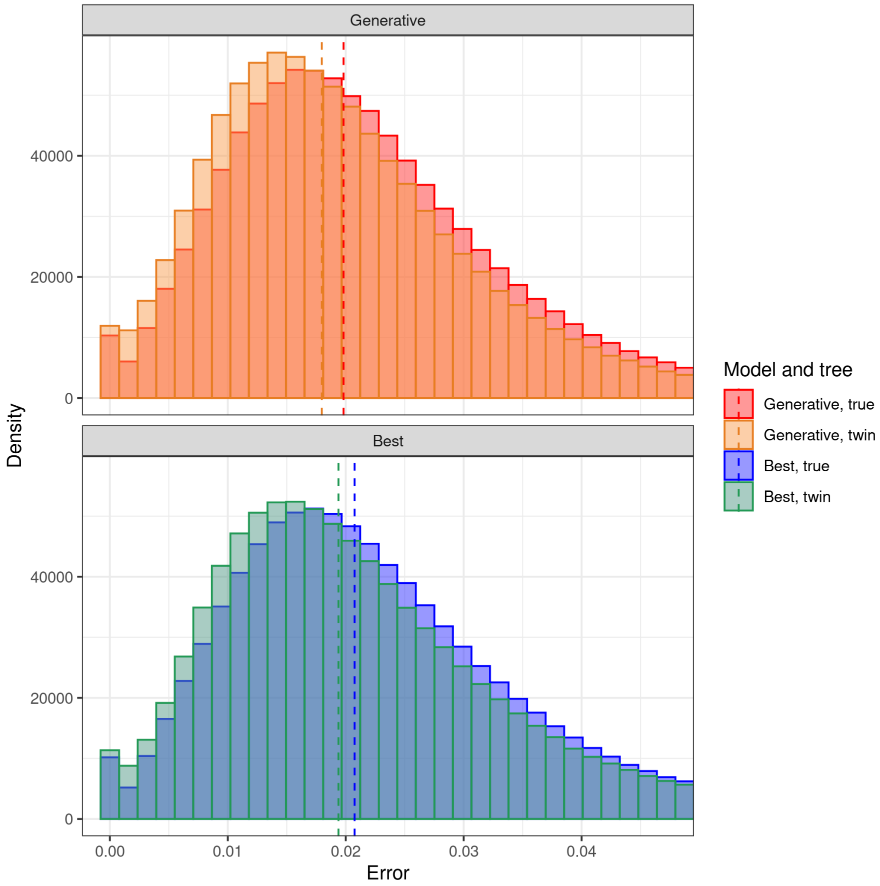


Figure 21: Aggregate error distributions for 100 replicates, for the tree distribution presented in 5.11 but with a per-nucleotide mutation rate of 0.25 / crown age. This took 1.8 days (wall clock time) to compute.

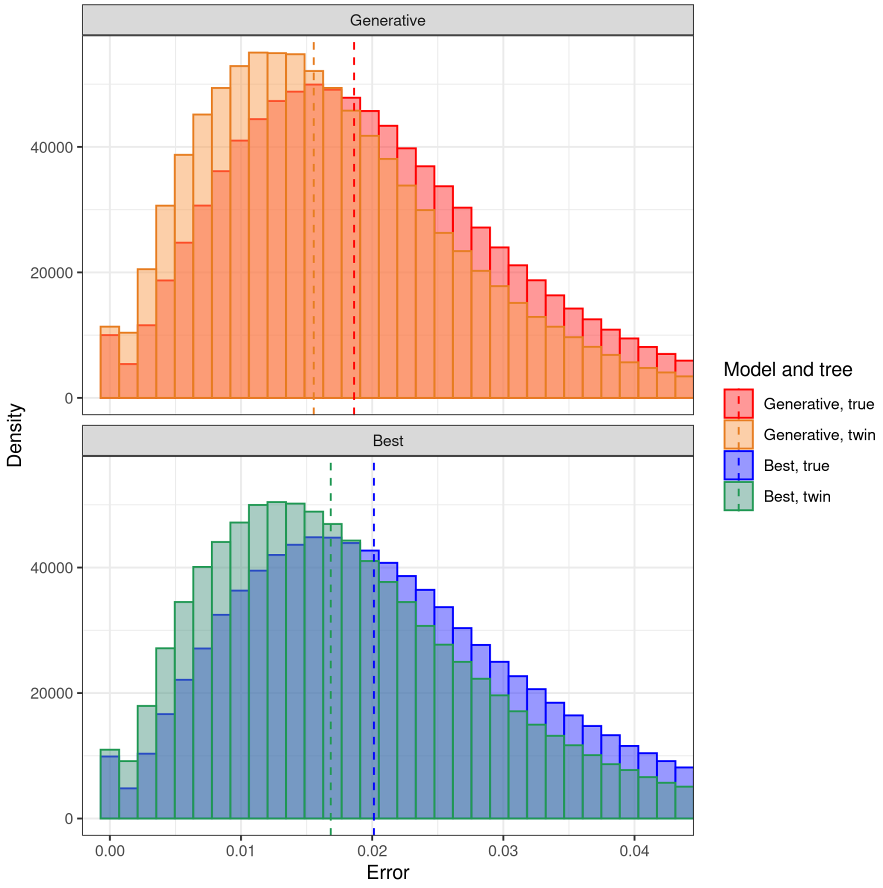


Figure 22: Aggregate error distributions for 100 replicates, for the tree distribution presented in 5.11 but with a per-nucleotide mutation rate of 0.50 / crown age. This took 2.2 days (wall clock time) to compute.

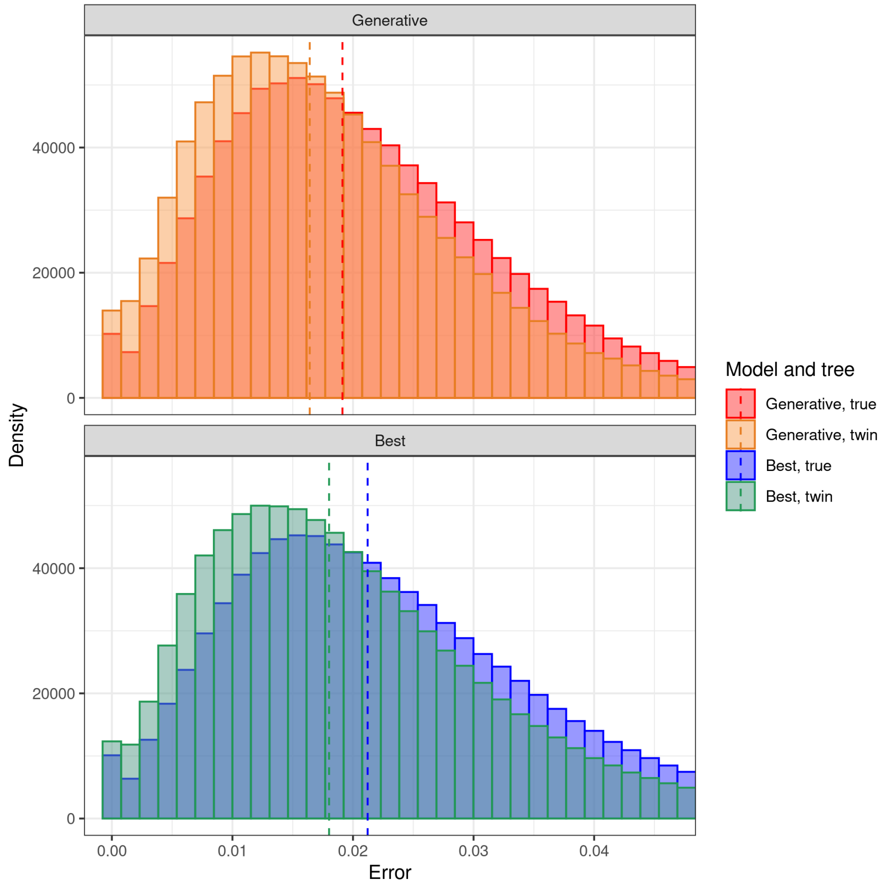


Figure 23: Aggregate error distributions for 100 replicates, for the tree distribution presented in 5.11 but with a per-nucleotide mutation rate of 0.75 / crown age. This took 2.5 days (wall clock time) to compute.

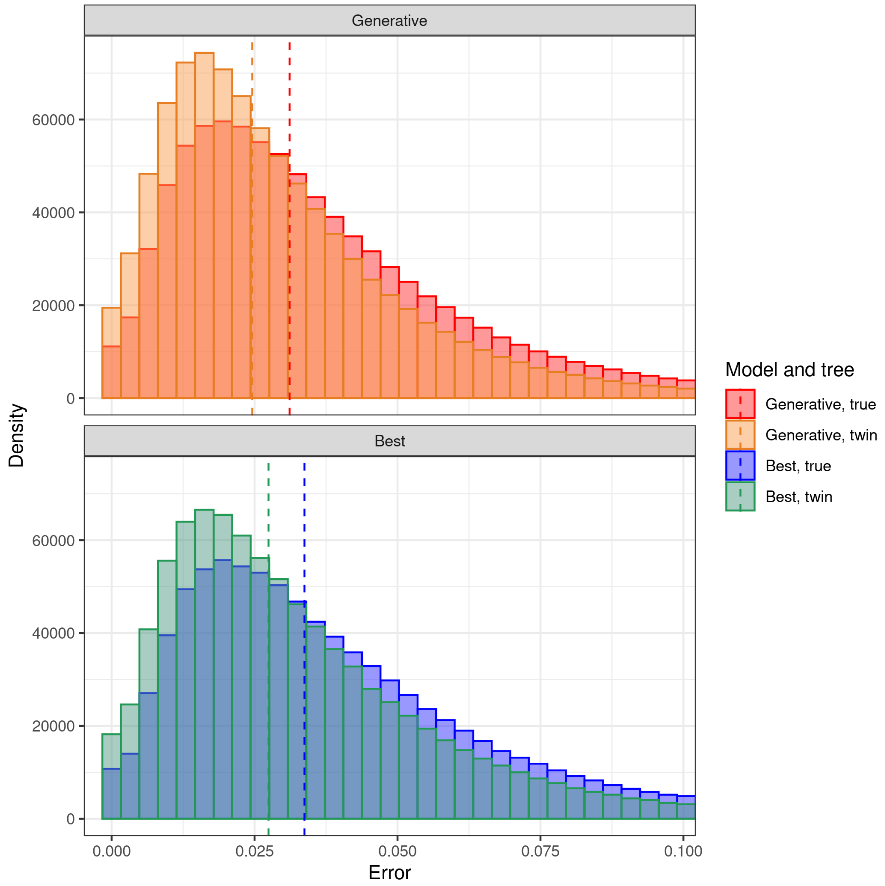


Figure 24: Aggregate error distributions for 100 replicates, for the tree distribution presented in 5.11 but with a per-nucleotide mutation rate of 1.25 / crown age. This took 2.9 days (wall clock time) to compute.

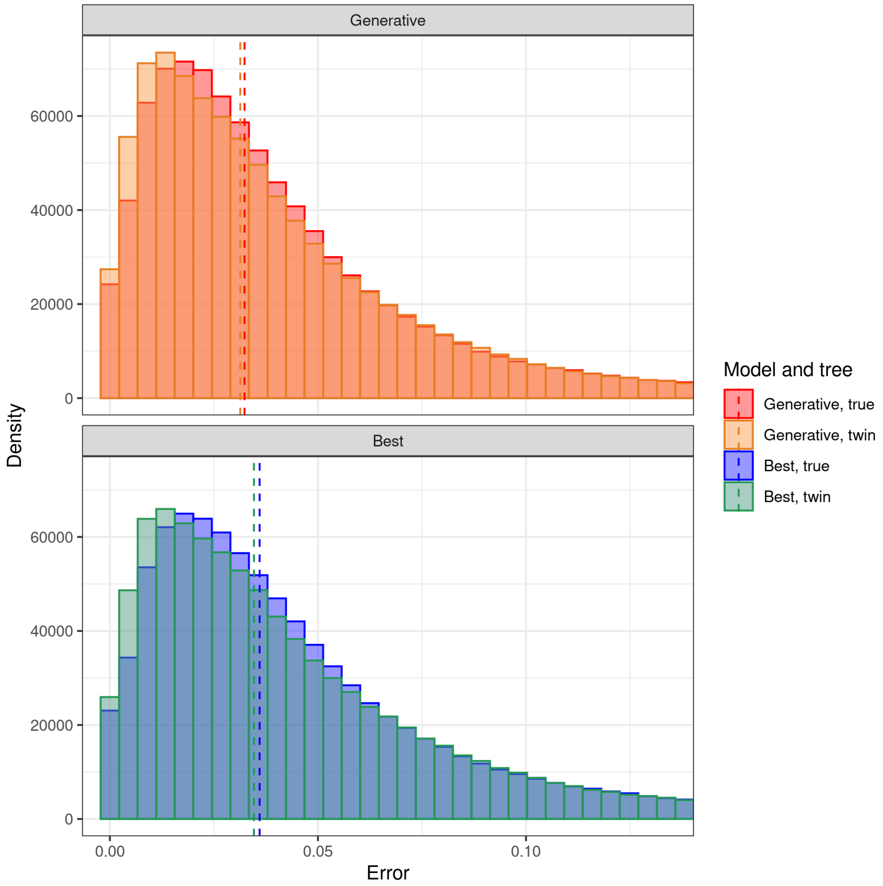


Figure 25: Aggregate error distributions for 100 replicates, for the tree distribution presented in 5.11 but with a per-nucleotide mutation rate of 1.50 / crown age. This took 3.0 days (wall clock time) to compute.

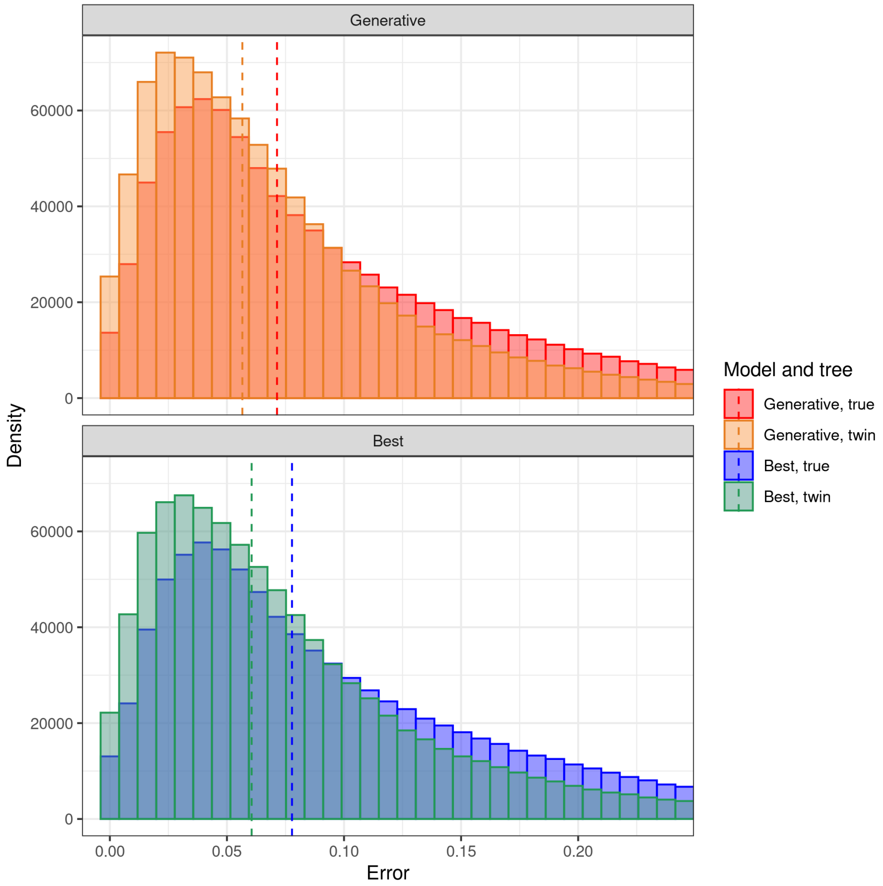


Figure 26: Aggregate error distributions for 100 replicates, for the tree distribution presented in 5.11 but with a per-nucleotide mutation rate of 2.0 / crown age. This is done for 100 replicates. This took 3.0 days (wall clock time) to compute.

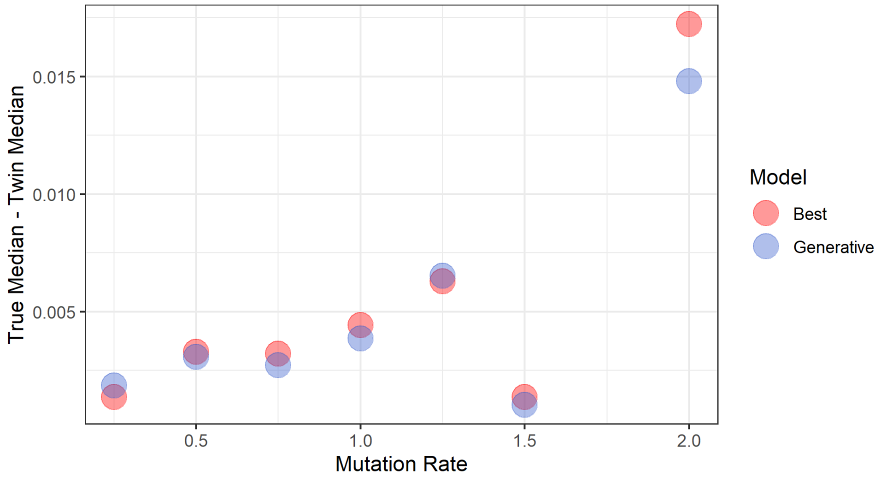


Figure 27: Difference between median true error and median twin error for different values of mutation rate.

### 6 Acknowledgments

We thank the Center for Information Technology of the University of Groningen for its support and for providing access to the Peregrine high performance computing cluster. We thank the Netherlands Organization for Scientific Research (NWO) for financial support through a VICI grant awarded to RSE.

### 7 Data accessibility

All code is archived at `http://github.com/richelbilderbeek/pirouette_article`, with DOI `https://doi.org/12.3456/zenodo.1234567`.

### 8 Author contributions

RJCB, GL and RSE conceived the idea for the package. RJCB created, tested and revised the package. GL provided major contributions to the package. RJCB wrote the first draft of the manuscript, GL and RSE contributed to revisions.
